# D155Y Substitution of SARS-CoV-2 ORF3a Weakens Binding with Caveolin-1

**DOI:** 10.1101/2021.03.26.437194

**Authors:** Suchetana Gupta, Ditipriya Mallick, Kumarjeet Banerjee, Shrimon Mukherjee, Soumyadev Sarkar, Sonny TM Lee, Partha Basuchowdhuri, Siddhartha S Jana

## Abstract

The clinical manifestation of the recent pandemic COVID-19, caused by the novel SARS-CoV-2 virus, varies from mild to severe respiratory illness. Although environmental, demographic and co-morbidity factors have an impact on the severity of the disease, contribution of the mutations in each of the viral genes towards the degree of severity needs a deeper understanding for designing a better therapeutic approach against COVID-19. Open Reading Frame-3a (ORF3a) protein has been found to be mutated at several positions. In this work, we have studied the effect of one of the most frequently occurring mutants, D155Y of ORF3a protein, found in Indian COVID-19 patients. Using computational simulations we demonstrated that the substitution at 155th changed the amino acids involved in salt bridge formation, hydrogen-bond occupancy, interactome clusters, and the stability of the protein compared with the other substitutions found in Indian patients. Protein-protein docking using HADDOCK analysis revealed that substitution D155Y weakened the binding affinity of ORF3a with caveolin-1 compared with the other substitutions, suggesting its importance in the overall stability of ORF3a-caveolin-1 complex, which may modulate the virulence property of SARS-CoV-2.

## 1. Introduction

The Severe Acute Respiratory Syndrome Coronavirus 2 (SARS-CoV-2) is the causative agent of the novel Coronavirus Disease 2019 (COVID-19) (1). Till August 4, 2021, 204.1 million cases have been reported worldwide spanning across 220 countries and territories, out of which 31.9 million people have been infected with SARS-CoV-2 in India. Mortality rate across the world varies drastically from 9.3% (Peru) to 0.3% (UAE) (2; 3). Although age, ethnicity and sex contribute to the demographic variation in the viral transmission and its case fatality rate, how mutation in viral genome, can change in such variation for pathological manifestation needs to be explored. The SARS-CoV-2 genome consists of approximately 30 kilobases and shares about 82% sequence identity with both SARS-CoV and MERS-CoV. It also shares more than 90% sequence identity for essential enzymes and structural proteins (4). Despite the similarity, only SARS-CoV-2 shows severe pathological manifestations in humans, suggesting the existence of differential molecular interactions between viral proteins and host cell machinery in COVID-19.

The SARS-CoV-2 genome broadly consists of 14 open reading frames (ORF), which are generated from nested transcription of subgenomic RNAs. With an exception, ORF1a and 1b encode for 16 non-structural proteins (nsp) known as replicase/transcriptase complex. The other ORFs code for 4 structural proteins and 8 accessory proteins (1; 4). ORF3 is wedged between spike (S) and envelop (E) ORFs and encodes for a membrane-spanning, ion channel protein ORF3a. It is also known as the single largest accessory protein of 275 amino acids (5). Ribosomal profiling has identified two putative overlapping genes, namely ORF3b and ORF3c, at the 3’ end of ORF3 with an alternative reading frame to the canonical ORF3a (6; 7; 8), whose functional importance is not well understood. ORF3a can localise at plasma membrane and Golgi complex, and can exist in both glycosylated and non-glycosylated forms (9). This viral protein has been shown to be highly immunogenic as antisera isolated from SARS-CoV-infected patients can detect ORF3a (9). Yount et.al and others have shown that ORF3a has been co-evolved with Spike (S) protein, suggesting the possibility of direct or indirect interactions between ORF3a and S protein (10; 11; 12). Studies in SARS-CoV-infected Caco2 cells show that ORF3a can also be efficiently released in detergent-resistant membrane structures and the diacidic motif, ExD, located within the domain VI and can play important role in membrane co-localisation (13). ORF3a has multi-functional roles including activating NLRP3 inflammasome and NFkB pathway, upregulating fibrinogen secretion, downregulating IFN Type I and inducing ER stress and pro-apoptotic activity (5; 14; 15; 16). Therefore, mutations in this protein warrant further study to understand their role in the virulence and immune evasive potential of the recent SARS-CoV-2. Several mutations have been reported in the ORF3a gene and have been classified in the form of clades and sub-clades. The mutation patterns of ORF3a gene have been characterized as largely non-synonymous and increasingly deleterious (Q57H, H93Y, R126T, L127I, W128L, L129F, W131C, D155Y, D173Y, G196V, and G251V). G251V and Q57H exhibit severe virulence property (17; 18; 19; 20).Viral infection consists of several steps starting from viral entry, intracellular trafficking, replication, assembly and then release of the viral particles. One of the host proteins like Caveolin-1 can not only interact with viral ORF3a but also regulate each of the processes in viral infection (21; 22; 23; 24). How the mutation at ORF3a can change the interaction between ORF3a and Caveolin 1 has not been clearly understood. Our study aims to understand the effect of one of the frequently occurring substitutions, D155Y, in the structural stability of the ORF3a protein and its ability to form complex with caveolin-1. Using computational simulation, protein-protein docking we find that the substitution of D with Y at 155th position causes instability of ORF3a-caveolin-1 complex.

## 2. Methods

### 2.1. Bioinformatics Methods

A total of 415,309 sequences of ORF3a protein deposited in NCBI database as on August 4, 2021 were considered for the bioinformatics analysis. The keywords used for the search were “SARS-CoV-2”, “ORF3a protein”, and “complete structure”. These structures were aligned using the BLAST algorithm on the NCBI website. Some of the post-BLAST sequences were larger than 275 due to erroneous performance of the code. But such cases were very low in number. Subsequently, the erroneous sequences were manually cleaned to obtain the final alignments of the complete protein sequences (275 amino acids). The total number of samples whose locations were geo-tagged to India was 1452. These sequences were compared with the Wuhan sequence (NCBI Accession No: *Y P* 009724391.1 (25)). The positions, where mismatches were observed with respect to the Wuhan sequence (WT), were considered as locations of mutations. Clearly, lesser number of mutations denote a sequence more similar to the WT, whereas more number of mutations denote a sequence more deviant from the WT. The sequences from NCBI database were compared with the WT and the number of mutations at every position of ORF3a were computed for further analysis. We calculated the percentage of occurrence (POC) of each mutation in Indian with respect to the world, using the following formula, POC = [No. of mutations at respective position (India)/ No. of mutations at respective position (World)] × 100. This essentially provides us with the frequency distribution of the mutations found at each position of ORF3a. We also calculated the substitution frequency (f) of D155Y and S171L worldwide using the formula, f = (No. of mutations at respective position/ No. of total sequences reported) × 100. It reflects the global distribution of a certain mutation. This essentially provides us with the frequency distribution of the mutations found at each position of ORF3a. We have used PROVEAN score to assess whether the effect of a mutation is deleterious or neutral. PROVEAN score of each mutation was determined using PROVEAN web server (26).

### 2.2. Preparation of structure of the ORF3a protein

The cryo-EM structure of WT ORF3a protein of SARS-CoV-2 was obtained from PDB (PDB ID: 6XDC (27)). The symmetry information present in the PDB file was used to convert the structure into the functional dimeric form using PDBe PISA server online (28). The residues on the second monomer have been numbered using “ ‘ “ throughout the manuscript. We considered the PDB structure of ORF3a, which has residues from 40^*th*^ to 238^*th*^ as no homologous structure of the protein was available. We introduced the necessary mutations (D155Y and S171L) by modelling the residues in Swiss PDB Viewer (29).

### 2.3. Molecular dynamics simulation

We performed classical Molecular Dynamics (MD) simulation in AMBER20 using AMBER ff14sb force field (30; 31). The missing hydrogen atoms in the protein structure were added by the LEaP module of AMBER20 package. The protein was then subjected to energy minimisation for 2000 steps using steepest descent and conjugate gradient algorithms. We then solvated the energy minimised structures using rectangular water boxes comprising of TIP3P water molecules (32). Particle mesh Ewald method was used to calculate the electrostatic interactions at a cut-off distance of 12Å. We performed initial minimisation and equilibration in order to avoid bad contacts. This was followed by equilibration using NVT ensemble at 300 K for about 500 ps. The systems were then equilibrated using NPT ensemble at 1 atm pressure for 1 ns. We considered 2fs as the time step throughout the minimisation equilibration-production. After the energy values and the density values converged, the systems were subjected to 100ns production runs using NPT ensemble at 300K and 1 atm pressure. The coordinates were saved after intervals of 2ps. We performed the analyses CPPTRAJ module of AMBER and visualizations were performed using VMD (33; 34). To predict the free energies of binding of ligands to the receptor, the two most commonly used methods are the *molecular mechanics generalized Born surface area* (MM-GBSA) and molecular mechanics PoissonBoltzmann surface area (MM-PBSA) methods. Even though the PoissonBoltzmann (PB) method is theoretically much more rigorous than the generalized Born (GB) method, both the methods are equally efficient to predict the correct binding affinities [(1-5)]. Here, we used the MM-GBSA method to calculate the relative free energies for binding of ORF3a to caveolin-1. For a given complex, the free energy of binding is calculated as,

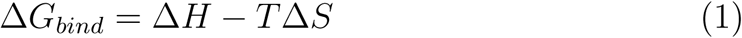

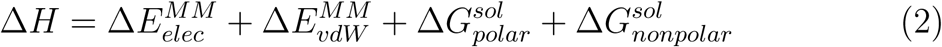

Where 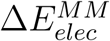 and 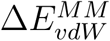 are the electrostatic and van der Waal’s contri-butions respectively, and 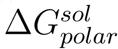 are the polar and nonpolarand 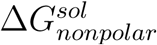 solvation terms, respectively. The nonpolar energy is calculated by the solvent accessible surface area (SASA) while the polar contribution is estimated using the GB model with an external dielectric constant of 80 and an internal dielectric constant of 4. We considered igb=5 for our calculations. As our calculations involve binding of similar types of ligands to the receptor,the entropic contributions are neglected. Thus, these computed values will be referred to as relative binding free energies. The binding energies for the complexes were calculated using MM-GBSA suite of AMBER (35).

### 2.4. Graph Theory

We used graph theory to decipher the composition of the interactomes in terms of participating amino acids involved in pairwise interactions. Briefly, the graph structure *G*(*V, E, W*) is denoted by three sets. The first one is, the set of residues, denoted as the vertex set (*V*) of a graph. Each individual residue, here, was considered as an independent entity, formally termed as a vertex or a node. Say, we denote *V* as {*v*_1_, *v*_2_, *v*_3_, …, *v*_*n*_}, where *v*_*i*_ is the *i*^*th*^ residue. Therefore, |*V*| = *n*, where *n* is the number of nodes in the graph, otherwise also known as the order of the graph. The second set is the set of interactions between residues denoted as the edge set (*E*). The interaction between the *i*^*th*^ and the *j*^*th*^ residue may be represented as an edge *e*(*v*_*i*_, *v*_*j*_) and the edge set *E* may be represented as {*e*_1_, *e*_2_, *e*_3_, …, *e*_*m*_}. Therefore, |*E*| = *m*, where *m* is the number of edges in the graph, otherwise also known as the size of the graph. Please note that the edges were not considered as directed because, there was no significance of the roles of the interacting residues in these interactions. We initially calculated the average energy values (calculated per unit time) over the time of observation for all ^*n*^*C*_2_ possible interactions. Some of them turned out to be high and were deemed insignificant. We used a threshold to remove the average energy values of those interactions. Here, *m* is the number of interactions with significant average energy values. These average energy values represented the importance of the interactions and were denoted as the set of edge weights (*W*). For every edge, there was a corresponding edge weight, therefore it could be concluded that |*W*| = *m*. In this work, we were interested in studying the interaction dynamics of the residues. Due to the difference in interaction energies, from observation, we could intuitively understand that a group of residues were more prone to interact among themselves than the other residues. But, to discover underlying densely interacting residue groups or clusters, we applied algorithms that could reveal the clusters accurately. The equivalent problem in residue-residue interaction graphs or networks is known as graph clustering. We have used one of the most popular community detection techniques, Louvain method, to find out the clusters in this residue interaction network (36). Please note that we have used the terms cluster and community interchangeably. It provided us with a cover *C* = {*c*_1_, *c*_2_, *c*_3_,….., *c*_*k*_}, where *k* is the number of communities and *c*_*i*_ is the *i*^*th*^ cluster/community. Each vertex in *V* belonged to exactly one of the clusters. Therefore, union of the vertex sets of all the clusters would lead back to *V*.

### 2.5. Modelling the protein-protein interaction complex

We used hierarchical approach to predict the structure of the protein in the absence of a suitable template structure for caveolin-1. I-TASSER server was used to generate five initial models (37; 38; 39). One model was selected based on the C-score (confidence score). The model was then evaluated using the SAVES v5.0 server, where Ramachandran plot and ERRAT analyses were performed (40; 41). Model visualizations were done using Chimera (42). This model was then simulated for 100ns to generate a more stable structure. The average structure was then considered as the initial structure for docking after proper structural evaluation by Ramachandran Plot and ERRAT analyses. The two molecules of human caveolin-1 were docked to the WT ORF3a by using HADDOCK (43). HADDOCK not only considers traditional energetics and shape complementarity, but also incorporates experimental data in terms of restraints to guide the docking of two proteins. The residues of domain IV of ORF3a and the residues Asp82 to Arg101 of human caveolin-1 were defined as active residues in docking based on the cryo-EM structure information (44; 45). On the basis of the most negative binding energy, we selected a starting structure for the WT ORF3a caveolin-1 complex. Similarly, we have considered the average simulated structures of D155Y and S171L variants of ORF3a and used them to dock to the human caveolin-1 protein via HADDOCK webserver. We generated a few complexes and considered the structures with the highest HADDOCK score as our starting complexes. We performed 100ns atomic MD simulations for all these complexes and analysed the data.

## 3. Results

### 3.1. Worldwide prevalence of D155Y substitution of ORF3a

ORF3a protein is important for the viral infection, spreading and modulating the host immune system. To understand the role of mutations in the function of this protein, we first checked the prevalence of each mutations found in ORF3a (Table S1). Fig. 1a shows the percentage of occurrence of mutant samples at each position of the ORF3a protein in Indian population in comparison to the global population affected by COVID-19 from a total of 415,309 samples, deposited in NCBI dataset (dated August 4, 2021). From this figure, we observe that mutations occur at 273 positions of the protein for the global population. Whereas, for the Indian population, mutations were found only at 94 positions. Although the number of instances for the mutation at the 57th position was the highest in both the global and the Indian population, we calculated the percentage of occurrence (POC) of each mutation in India. Mutations with highest POC values were further filtered by considering a threshold value of n=1000 counts for the world population and n=1 count for Indian population. We found 26 such mutations, which were shown in Fig 1b. We selected the positions 155th and 171st from the top-ranked (POC ≥ 1) and the middle-ranked groups (1>POC ≥ 0.1), respectively, for our study. The positions 155th and 171st showed 33 and 12 instances of mutations respectively in India, while the number of instances were 2,261 and 3,147, respectively, in the global population (Fig 1c). Note that the frequency of D155Y and S171L substitutions in India were 2.27 and 0.82, which were comparable to Asia.

**Fig. 1.**
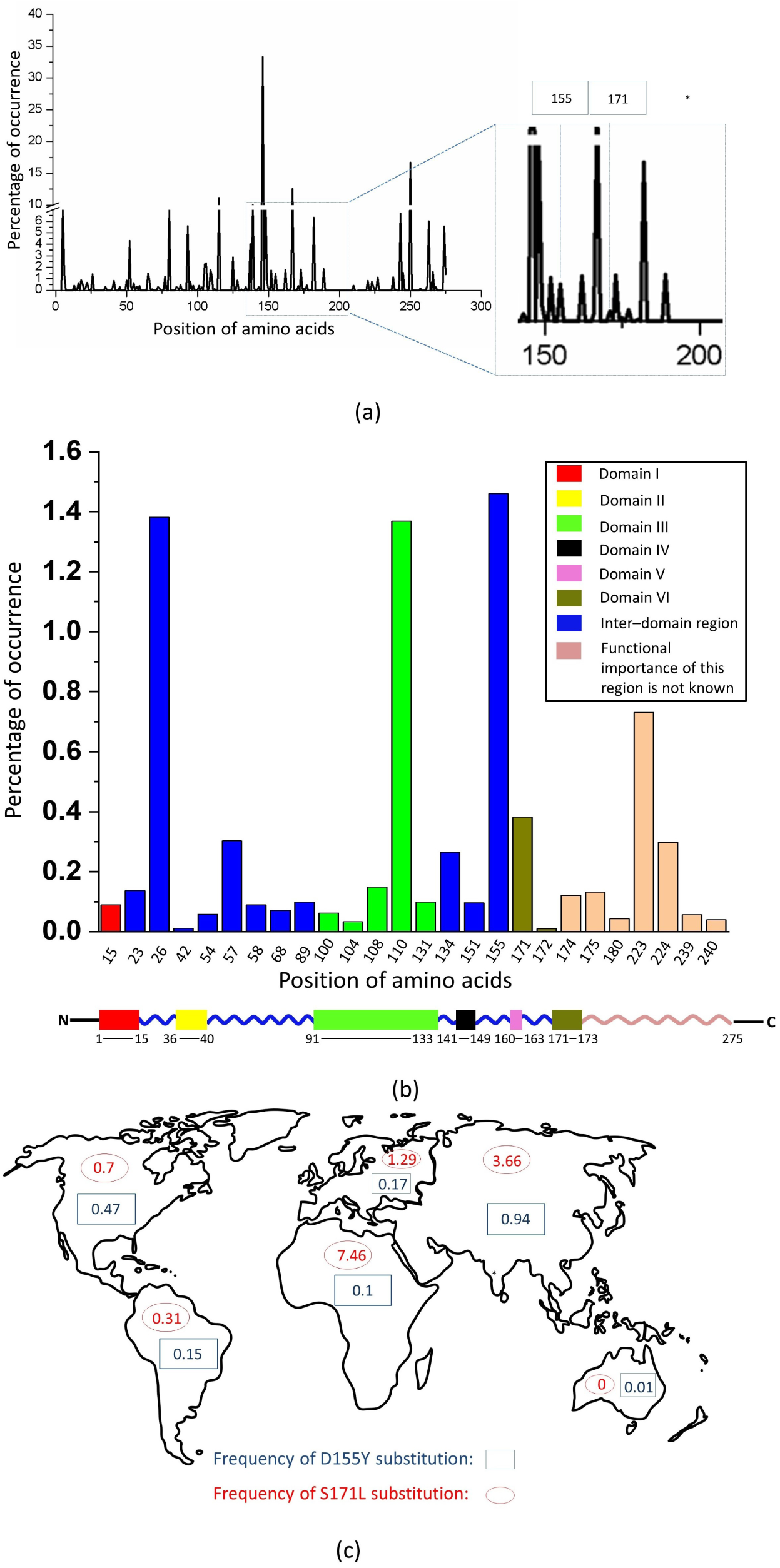
Distribution of mutations in ORF3a protein. Percentage of occurrence (*POC*) of various mutations at different positions of ORF3a protein found in India with respect to the world has been plotted in(a). 26 mutations have been arranged in descending order in terms of their *POC* values and were filtered on the basis of a pair of thresholds (threshold: n=1000 for the world and n=1 for India, n - No. of occurrence) and were plotted in (b). Color coded schematic representation of various domains of ORF3a. Lengths of the domains are not as per the scale. The global distribution of D155Y and S171L substitutions across the continents have been shown in blue and red respectively in (c). Percentage of occurrence of mutations in India with respect to the world (*POC*) is calculated using the formula: *POC*=[No. of mutations at respective position (India)/No. of mutations at respective position (World)] x100 and substitution frequency (*f*) is calculated using the formula, *f* =(No. of mutations at respective position/No. of total sequences reported) x 100.

### 3.2. Description of the protein system

The SARS-CoV-2 ORF3a protein can form dimer (46). The monomeric ORF3a has been divided into six domains, each having its own functional importance (19). Fig. 2 shows the locations of D155Y and D155’Y (red spheres) and S171L and S171’L (cyan spheres). The locations of these substitutions between domains IV and V, and in domain VI suggest their possible role in caveolin binding, intracellular protein sorting and intracellular membrane trafficking of ORF3a (19).

**Fig. 2.**
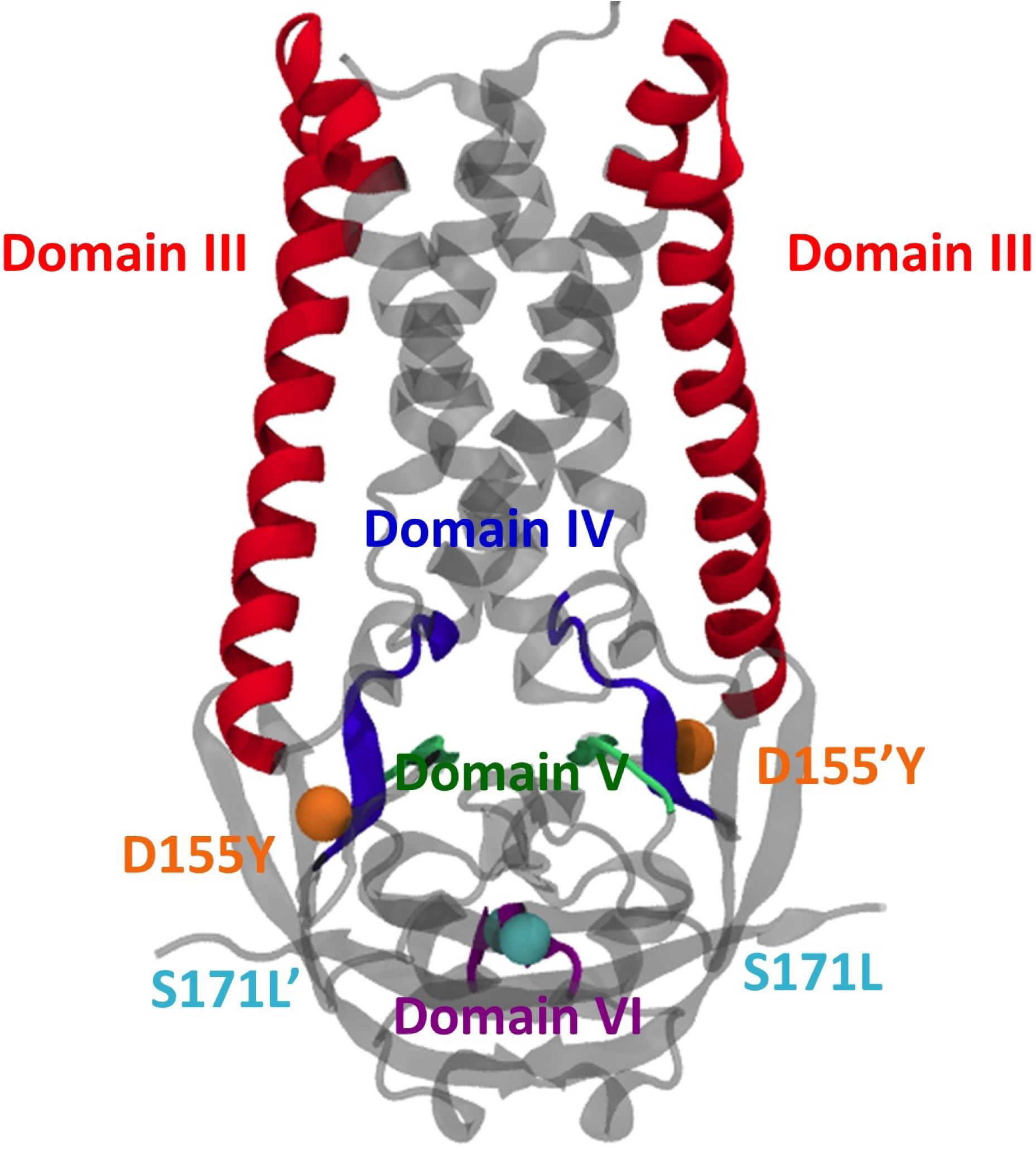
Structure of ORF3a protein. The structure of WT ORF3a (PDB ID: 6XDC) marking the functional domains as known from literature has been shown. The positions of mutation at the 155^*th*^ and 171^*st*^ positions have been shown in orange and cyan spheres respectively. Note that the 171^*st*^ residue of the second monomer of ORF3a (S171’L) is located on the posterior side of the image.

### 3.3. Stability of the two ORF3a variants, D155Y and S171L

The cryo-EM structure of WT ORF3a protein (residues 40 to 238) was downloaded from the Protein Data Bank (PDB ID: 6XDC) for analyzing the stability of ORF3a variants. The substitutions D155Y and S171L on each monomer were modelled on the WT structure separately using Swiss PDB Viewer (29). Each of these structures was simulated in triplicate till 200ns. Fig. 3 shows the time evolution of the root mean square deviation (RMSD) of the simulated structure with respect to the starting frame of simulation. WT and D155Y (black and red profiles, respectively) showed lesser RMSD (the final RMSD being 2.25Å) and lesser fluctuation, whereas the S171L (green profile) variant showed higher RMSD (the final RMSD being 2.75Å). The overall fluctuation in RMSD was also greater in S171L compared to the WT and D155Y. This indicates that S171L substitution causes more deviation. However, the final RMSD values attained by the WT and the two mutants were comparable, indicating a similar final simulated structure. So it can be concluded that the substitution at D155Y or S171L does not cause a major conformational change of ORF3a from the WT.

**Fig. 3.**
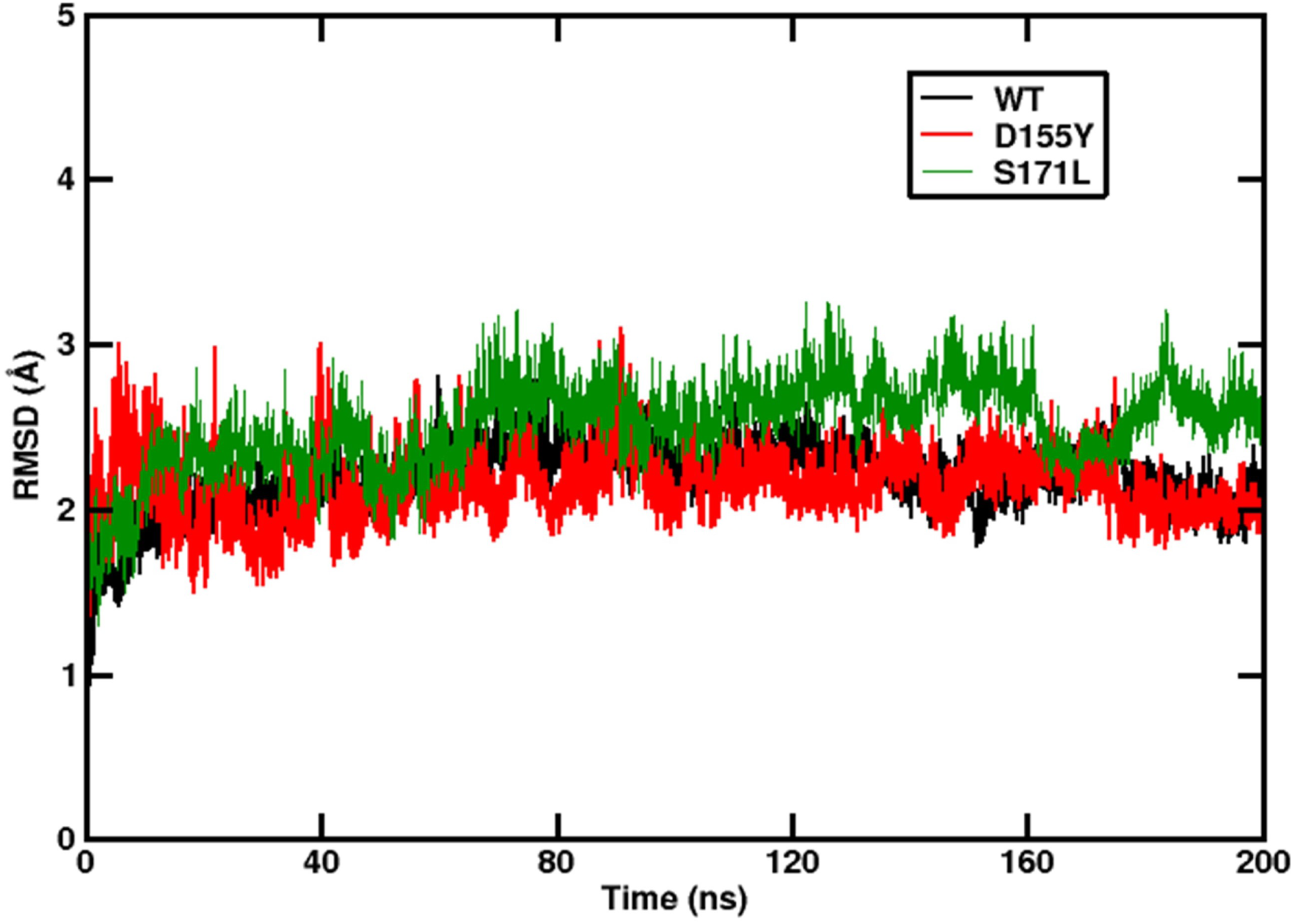
Stability of structure. The time evolution of the RMSD of the ORF3a proteins with respect to the starting structure. The data from the last 20ns of our simulation (stable trajectory) were considered and the snapshots were taken at intervals of 10ps for energy analysis. Production runs were repeated twice and the average of all the simulation sets were considered (Please see Fig S5). Black, red and green lines denote WT, D155Y and S171L, respectively.

### 3.4. Differential behaviour of the constituent residues

While RMSDs are a measure of the overall stability of the biological systems under consideration, the B-factor values give an idea on the flexibility of the individual residues. The B-factor values were measured to determine the average flexibility of the residues around their mean positions across all trajectories (Fig. 4). Fig. 4 shows that the residues in the WT ORF3a protein exhibit the least deviation from their mean position, whereas the residues in both mutants show more flexibility. Interestingly, in the D155Y variant, the 155^*th*^ residue showed higher flexibility compared to the WT (5.75 Å^2^ for WT, 9.12 Å^2^ for D155Y and 7.73 Å^2^ for WT, 10.60 Å^2^ for the D155Y at positions 155 and 155’ respectively). Similarly, in the S171L variant, the flexibility of the 171^*st*^ residue was higher than the WT (10.62 A^2^ for WT, 14.67 A^2^ for S171L and 22.06 Å ^2^ for WT, 26.97 Å^2^ for S171L at positions 171 and 171’ respectively). The terminal residues are exposed to solvent and are more flexible, resulting in their high B-factor values as seen in Fig. 4 (as marked). Several other residues, which are located both near the positions of mutations as well as distally also showed greater flexibility, suggesting an effect of the mutation on the overall dynamics of the protein at distant locations or mutations may have allosteric effects on the domain specific functions of the ORF3a protein.

**Fig. 4.**
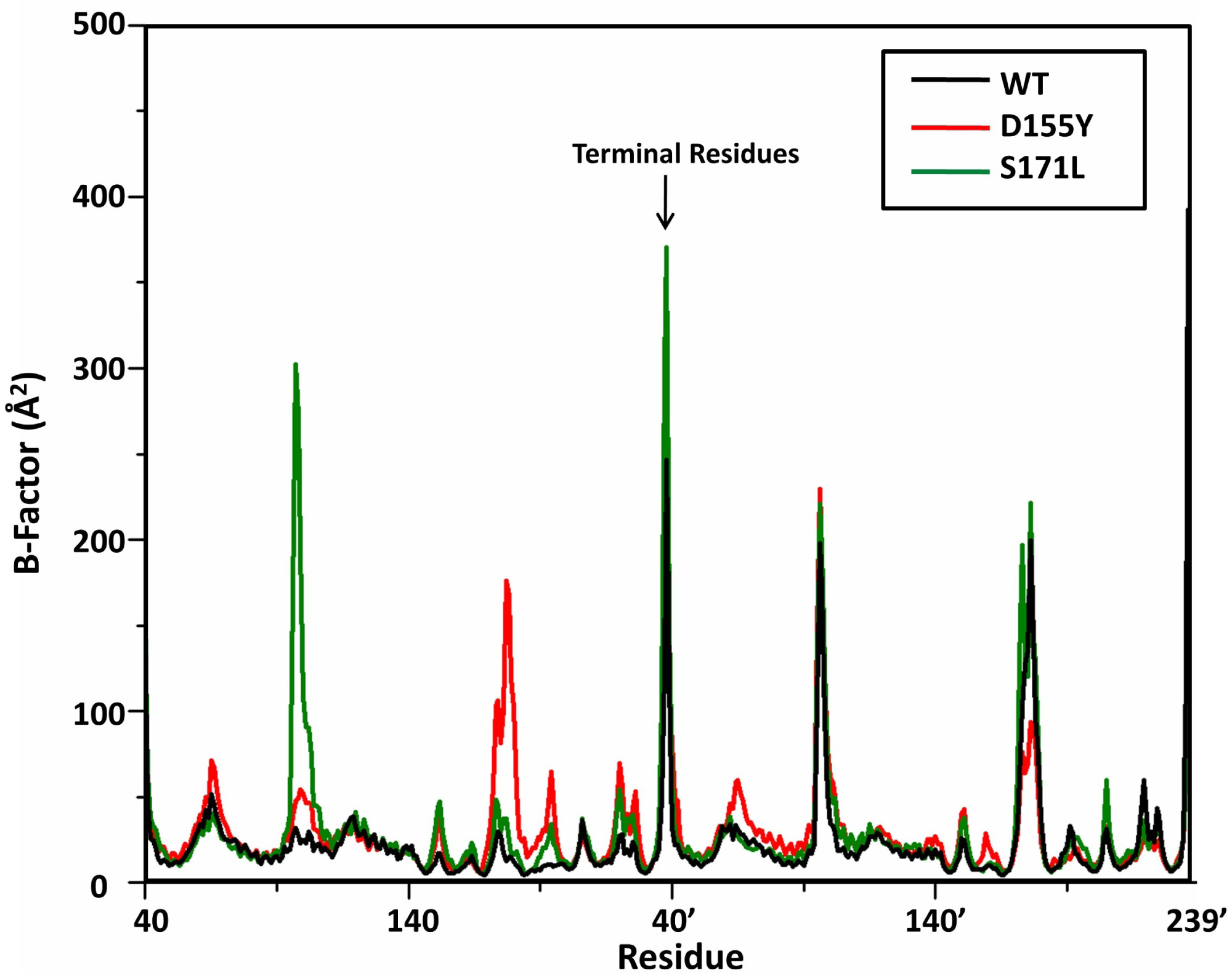
Flexibility of residues. The B-Factor plot for ORF3a for the three systems. Black, red and green lines denote WT, D155Y and S171L, respectively.

### 3.5. Differential contribution of stabilizing residue in the ORF3a proteins

The WT and the mutant ORF3a proteins were analysed to understand the role of individual amino acids, hydrogen bond occupancies and salt bridges in their structural stability. The free energies of the three variants (WT and the two mutants) were calculated by using the MM-GBSA module of Amber20 as tabulated in Table 1, which shows the differences in stability. The S171L mutant with a free energy of -5376.51 (± 19.34) kcal/mol was the most stable, followed by WT (−5356.85± 12.95kcal/mol) and D155Y mutant (−5266.41± 12.56kcal/mol). This indicates that the mutants D155Y and S171L can also exist independently just like the WT ORF3a protein. To understand the contributing factors for these variations in stabilizing energy, we looked at the contributions of each amino acid to the overall free energy and tabulated the top contributors for each variant in Table 2. While we observe that the group of residues contributing to the overall stability remains almost unchanged among the variants, their ranking differs. For instance, in WT, Arg68 plays the most important role, whereas in case of the mutant systems, it is Arg126’ that has the most contribution. However, Arg68 features as the second most contributory residue in D155Y mutant, whereas in the S171L mutant, it has the fourth position. In this mutant, Arg126 plays the second most important role. We also checked the hydrogen bond interactions in WT and the two mutant ORF3a proteins, and found that the total number of hydrogen bonds remain same (average number is 95, Fig.S1), in all the three variants. In contrast, the individual residues that have the most hydrogen bond occupancy vary among the ORF3a proteins. In WT, we found that the top three residue pairs involved in forming hydrogen bonds with the maximum occupancy are Tyr156’-Lys192’, Arg134-Asp155 and Leu203-Asp210. In D155Y, the top three residue pairs forming the hydrogen bonds with maximum occupancy are Tyr212’-Thr164’, Ser205’-Asn144’ and Ser205-Asn144. In the mutant S171L, the top three residue pairs forming hydrogen bonds with maximum occupancy are Leu203-Asp210, Thr89’-Leu85’ and Leu203’- Asp210’. A detailed list is given in Table S2. We also calculated the salt bridge interactions for the WT and the two mutant proteins and tabulated the list in Table S3, which shows that D155Y forms lesser number of salt bridges compared to the WT and the S171L (n=24 for D155Y and n=31 for WT and S171L). Interestingly, mutation at position 155, but not at 171, breaks the salt bridge formation between Asp155-Arg134. This residue pair is formed at the end of the alpha helix and the beginning of a beta sheet in the proximity of domains III and IV of ORF3a. The loss of salt bridge interaction in D155Y may play a significant role in the binding affinity of the interacting partner of the ORF3a protein at this region.

**Table 1:** The list of binding free energies for the three systems are given. The values in parentheses indicate their standard deviations.

| System | Free Energy (kcal/mol) |
| --- | --- |
| WT | -5360.30(12.95) |
| D155Y | -5263.66(12.56) |
| S171L | -5375.66(19.34) |

**Table 2:**
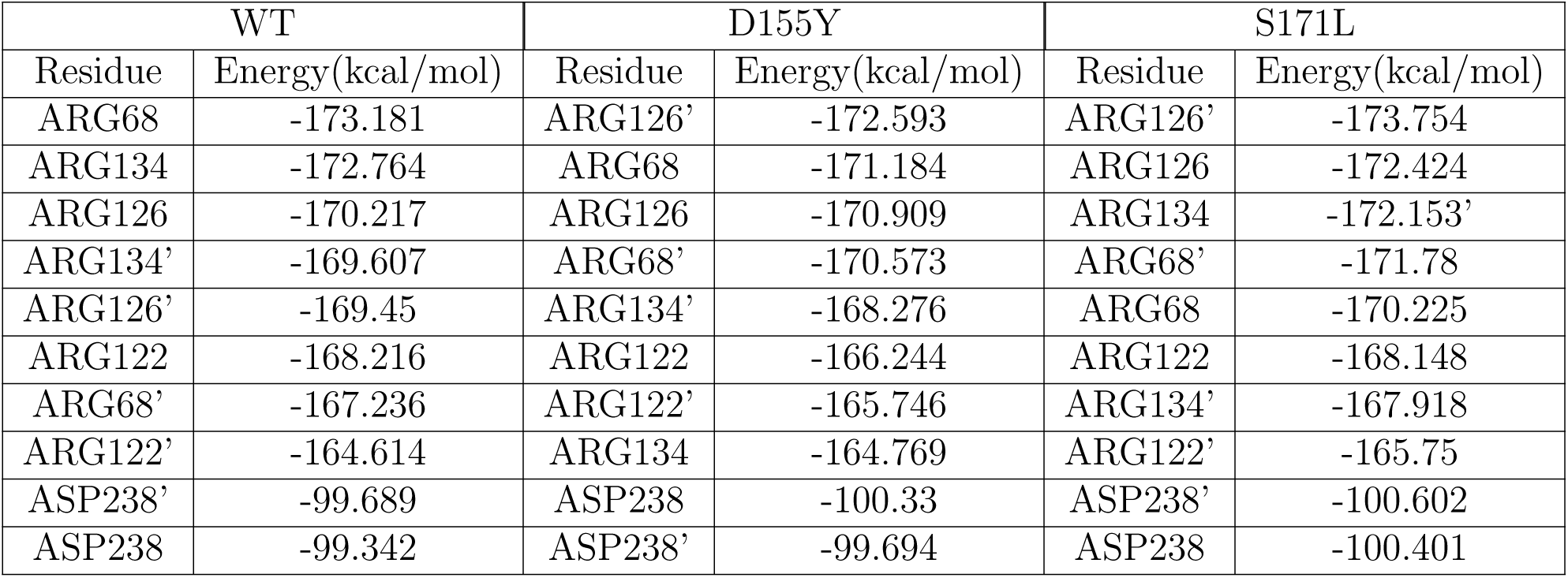
List of residues contributing to the overall stability of the systems.

**Table 3:** The list of binding free energies for the different mutant ORF3a proteinsare given. The values in parentheses indicate their standard deviations.

| System | Free Energy(kcal/mol) |
| --- | --- |
| Q57H | -5256.70 (69.71) |
| Q57H-D155Y | -5113.04 (43.07) |
| Q57H-S171L | -5251.63 (68.42) |
| W131C | -5364.58 (46.83) |
| W131R | -5740.51 (53.43) |
| G172C | -5305.72 (58.26) |
| G172V | -5379.19 (61.07) |

### 3.6. Changes in interactome interactions - A graph theoretic perspective

The variation in hydrogen base pairing and salt bridges prompted us to check the interactome interactions in ORF3a variants. We represented the interactions between the interactomes in terms of a network. The pairwise hydrophobic interaction energies of the residues in the WT and the two mutants of ORF3a were calculated using MMGBSA suite of Amber20. These hydrophobic interaction energies were considered for building a residue-residue interaction network for the WT and the two mutant proteins. Here, we have used graph data structures and relevant algorithms, to model the interactions among the residues. The spatial orientation of the protein, adjacency of the residues and interactions among them play a role in finding the clusters or communities of interacting residues. In the visualization of the clusters, as seen in Fig. 5a, we see the whole interaction network and an overview of the clusters. In Fig. 5b, we zoom on one part of the graph and provide a closer view of the interactions. The node colours denote its affiliation to a certain cluster. The edge colour is determined by the colours of the nodes it is incident upon. The edge thickness denotes the strength of the interactions between the residues, i.e., the weight of the edge. In Fig. 5c, one residue has been selected to show the nodes adjacent to it (also known as its neighbourhood). Fig. 5a shows the spatial orientation of the clusters, which can be seen in the actual protein. We observe that the membership of the residues in the clusters in each protein has shown substantial variation. A list of the clusters and their constituent residues has been provided for the WT and the two mutants (D155Y and S171L) in Table S4. Fig. 6 and Table S5 indicate that the residues of the functional domains have rearranged in different interacting clusters in WT and the two mutants. Domain III being the largest in size has split into the most number of clusters. However, the clusters are different in terms of the constituent residues for WT and the two mutants. Thus, we may conclude that the mutations have changed the interaction patterns of the interactomes present in the protein. Due to changes in residue interactions, the clusters have changed from WT to the other mutants. But it should be noted that the cluster membership for the nodes in the regions of mutations do not change. This indicates that the mutations have distal effects too, which can be explored further for better understanding of the protein function.

**Fig. 5.**
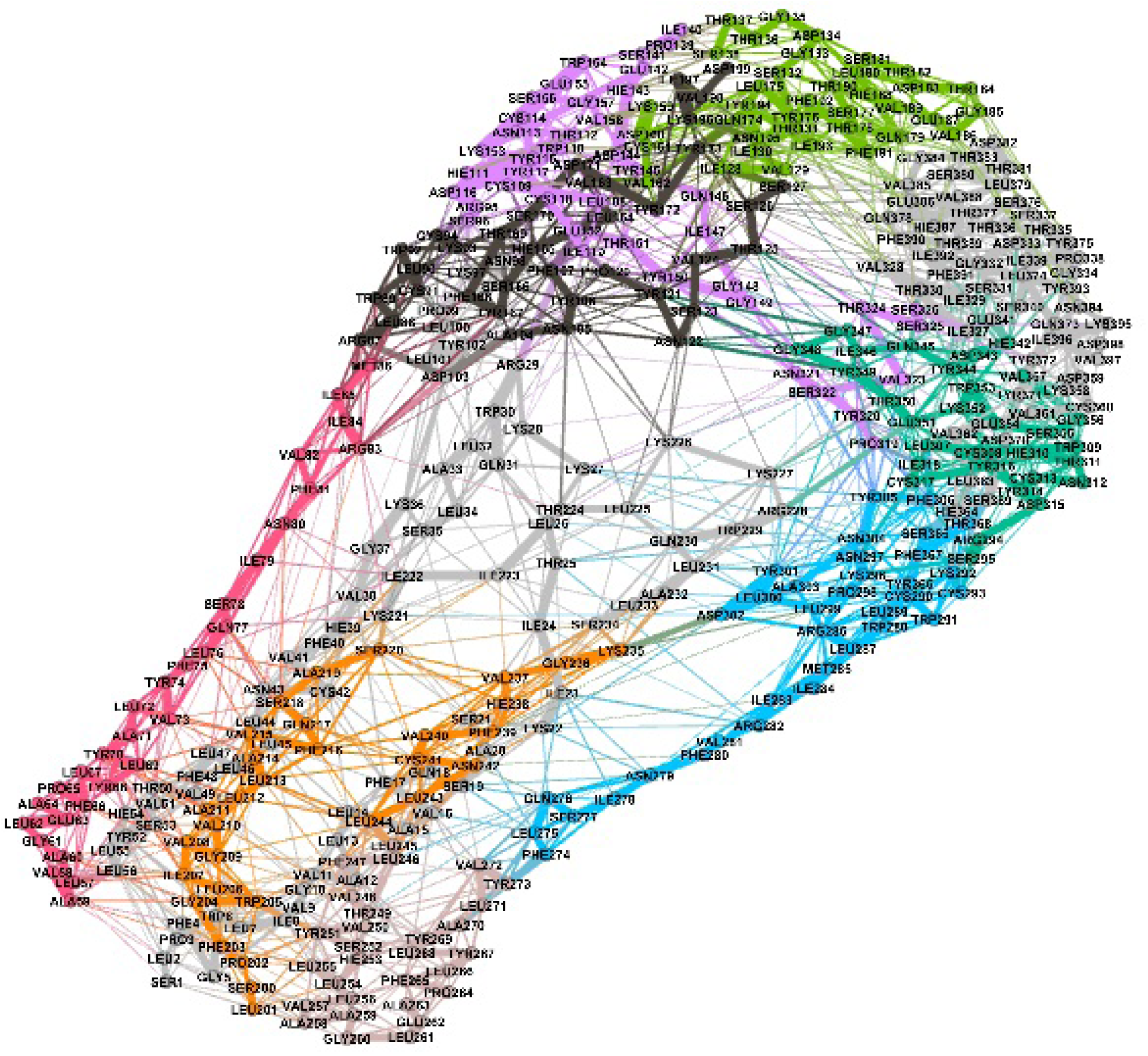
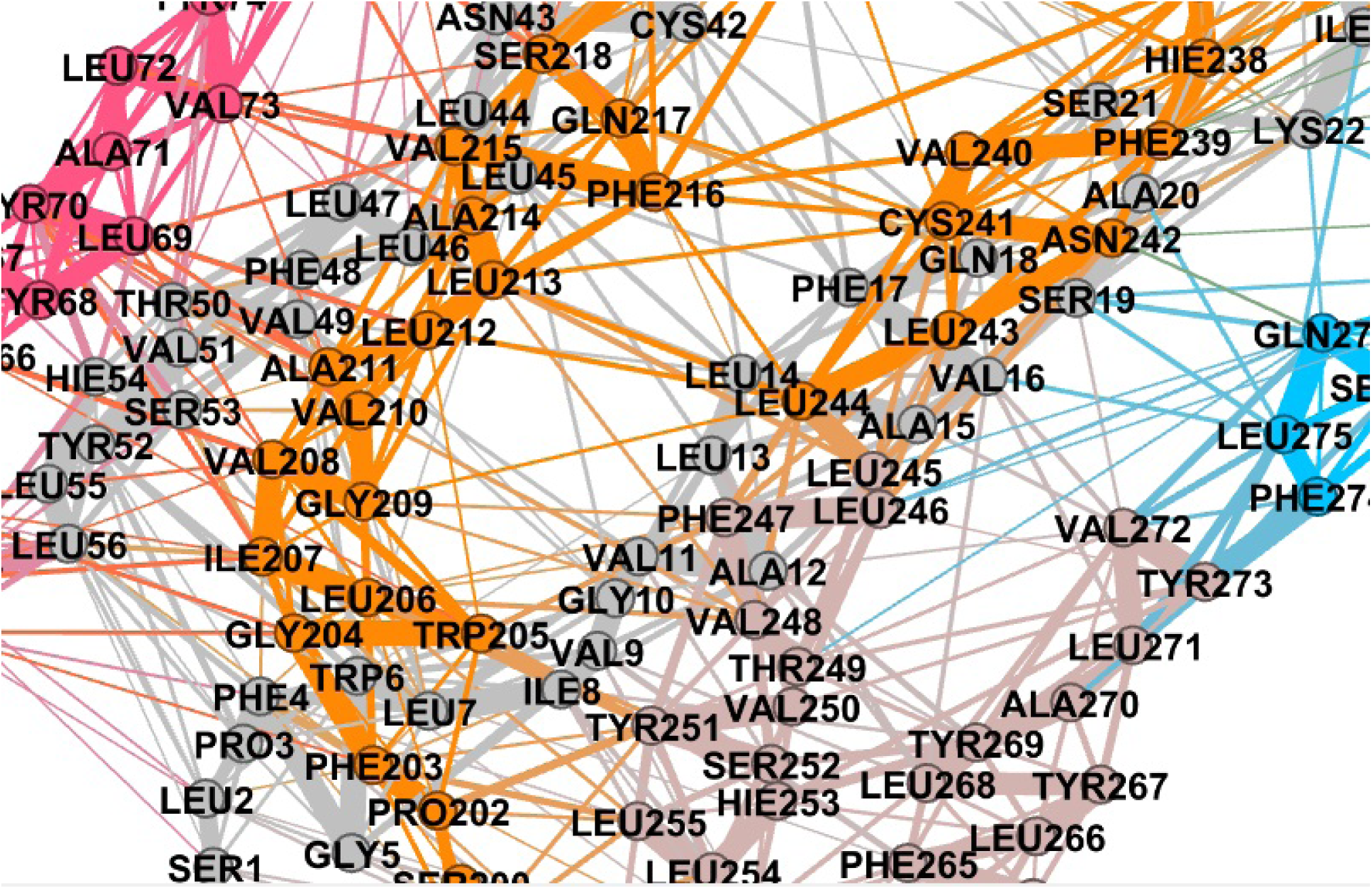
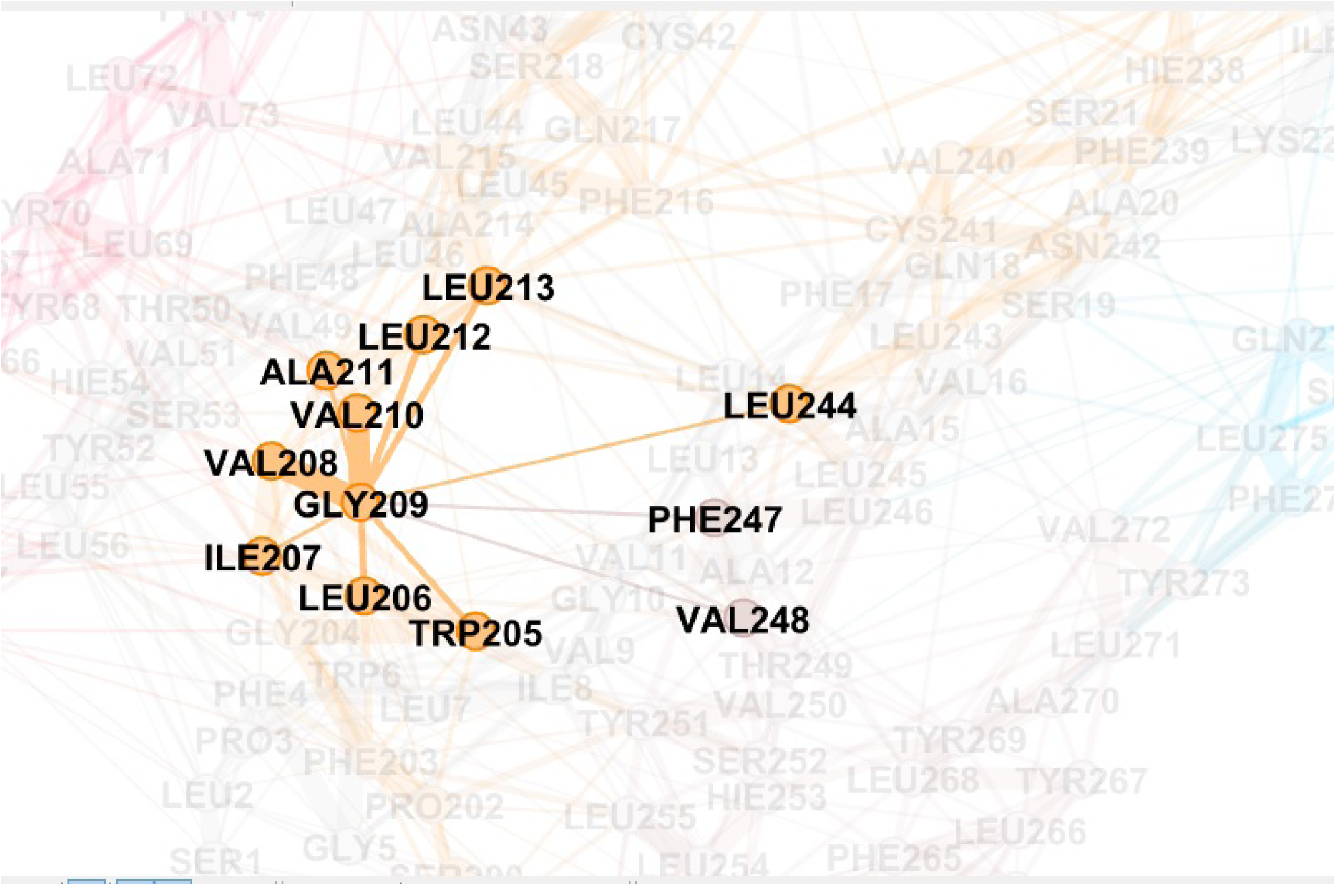
Visualization of residue interaction network in WT ORF3a protein using Gephi (59). (a) The whole residue interaction network showing the complete cover C, with nodes coloured with the membership colour of a particular cluster, (b) A magnified view of the residue interaction network, and (c) Shows one particular residue (here, GLY209) and the residues it is directly interacting with.

**Fig. 6.**
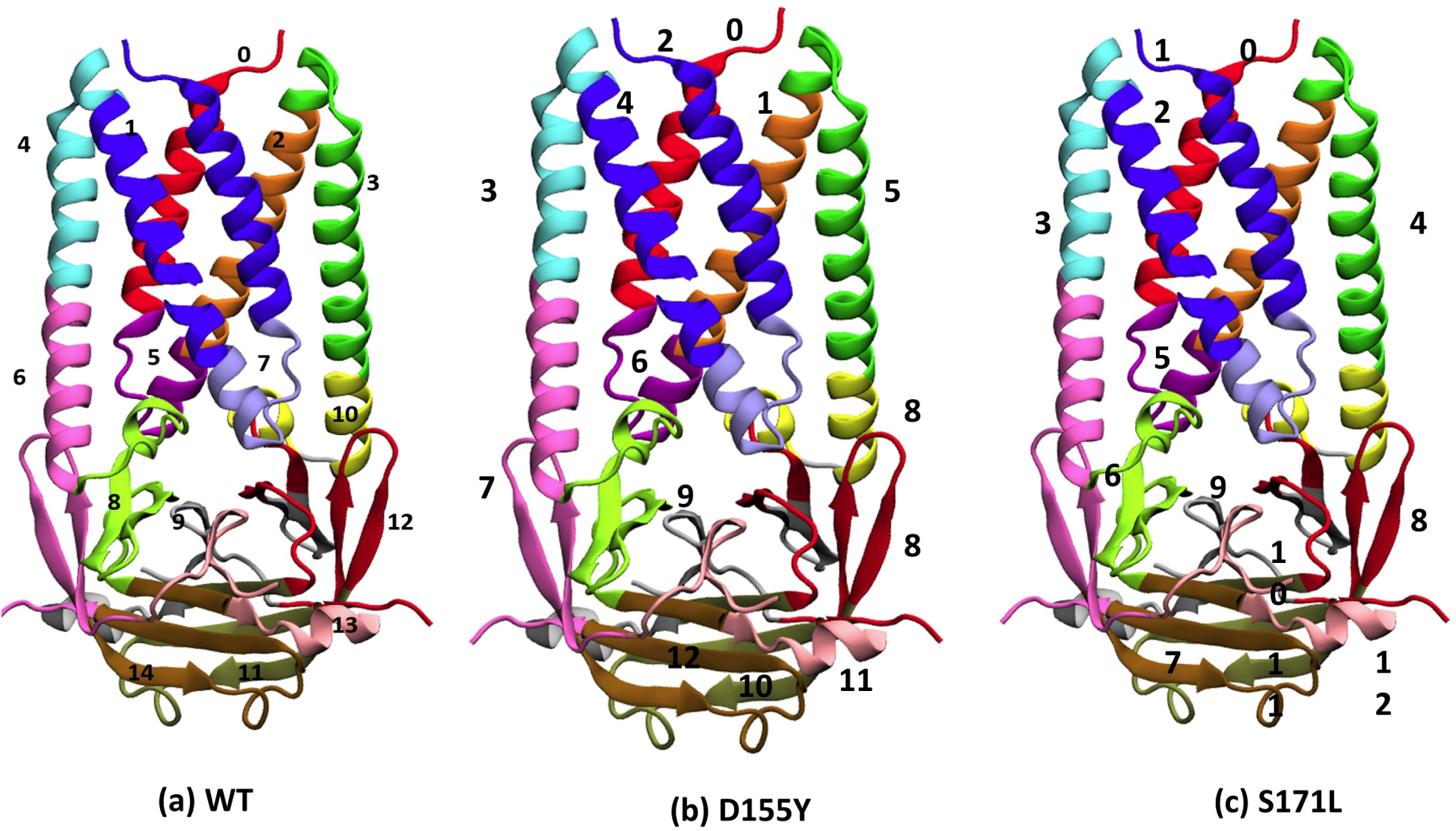
Interactome clusters in SARS-CoV-2 ORF3a protein. The different interactomes are shown for (a) WT, (b) D155Y and (c) S171L. fThe list of residues constituting each cluster for WT and the two mutants has been numbered as 0 to 14 (for WT ORF3a) and 0 to 12 (for mutants). The positions of the cluster numbers as mentioned in Table S4. have been indicated.

### 3.7. Formation of complex with the partner protein caveolin-1

The 155^*th*^ residue of ORF3a resides in between domains IV and V. It has been shown that domain IV of ORF3a binds with caveolin-1 protein. Therefore, we are interested to check if the substitution at this position can change the binding interaction of ORF3a protein with host caveolin-1. Issa et al. and others have suggested that interaction between ORF3a and caveolin-1 is required for viral uptake and regulation of viral life cycle (19; 23; 45). We modelled caveolin-1 using a hierarchical approach to predict the structure of the protein. Five initial models were then generated using the I-TASSER server. Out of these, one model was selected based on the C-score (confidence score). This model was then evaluated using the SAVES v5.0 server, where Ramachandran plot analysis and ERRAT analysis were performed as shown in Fig. 7a-b. Ramachandran plot showed that 96.9% of the residues of caveolin-1 were within the favoured and allowed regions, while 3.1% of the residues were in the disallowed regions. On the other hand, ERRAT analysis had an overall quality factor of 89.412. The modelled structure of human caveolin-1 was simulated for 100ns to generate a well equilibrated and stable structure. The stability of the simulation, as evident from the time evolution of the RMSD of the protein from its starting structure, has been shown in Fig. S2a. The average structure from this simulation (Fig. S2b) was considered as the starting structure of the ORF3a-caveolin-1 complexes after proper structural evaluation. Ramachandran plot of the average simulated and stable structure showed that 98.1% of the residues were within the ranges of favourable and allowed regions, and only 1.9% of the residues were in the disallowed regions (Fig. 7c-d). ERRAT plot too showed an improvement with the overall quality factor increased to 94.304%.Thus, both Ramachandran plot and ERRAT analysis (Fig. 7c-d) indicated that the caveolin-1 model was of acceptable quality, and could be used as the starting structure for docking. Similarly, we also carried out ERRAT analysis and Ramachandran plot for the starting structure (PDB ID: 6XDC) and average simulated structure of ORF3a (as shown in Fig. S7). ERRAT overall quality factor increased from 92.988 in the initial structure to 96.409 for the average simulated structure. For the Ramachandran plot, residues in the most favored regions increased from 89.4% to 93.4%. We carried out our protein-protein docking using the HADDOCK webserver (43; 47). We consider binding domains on ORF3a and caveolin-1 as the interacting residues (44; 45). Our analysis generated twelve probable structures from three clusters as shown in Fig. S3. In each of these structures, we had two molecules of the human caveolin-1 interacting with the dimeric form of ORF3a protein. The first structure from the cluster 1 (left-most structure in first row of Fig. S3), having a HADDOCK score of -155.3 (±22.2) was considered as our starting structure. This structure showed a symmetrical nature. The buried surface area of this complex was found to be 3031.4 (± Å^2^181) A, signifying a strong complex. We used the H++ webserver to check for protonation state of the target protein, caveolin. Since we considered pH 7, there was no change in the protonation state of the target protein^*^. The starting structures for WT and the two mutants were simulated for 100ns. The stability for these structures was assessed by plotting the time evolution of their RMSD values with respect to the starting frame of simulation (as shown in Fig. 8). We observed that the WT and the S171L are stable having an average RMSD value of around 15Å. Although the absolute value is high, yet the protein complexes reached stability and showed a plateau in the RMSD plot from 40ns, again indicating a stable complex. However, for the D155Y system, the protein complex showed a lot more fluctuation and deviation from the starting structure. This indicates a not-so-stable complex structure, which is further supported by the lower PROVEAN score of D155Y (Table S6). Since the mutation is present in the vicinity of the caveolin binding domain in ORF3a, it can be said that the presence of the mutation leads to an unstable protein-protein complex formation. Thus, the D155Y substitution interferes with the caveolin binding activity of ORF3a protein. We also calculated the free energy, corresponding to the binding of caveolin-1 to the ORF3a protein, in these three protein complex systems. The values for WT, D155Y and S171L were -37.64 (±8.32) kcal/mol, -20.31 (±8.60) kcal/mol and -28.39 (±0.71) kcal/mol, respectively. From these values, it is evident that the binding affinity of caveolin-1 is considerably less in D155Y mutant compared to WT and S171L. This is in corroboration with the unstable protein protein complex in the D155Y system. The change in hydrogen bonding, salt bridge pattern and hydrophobic interaction pattern associated with D155Y substitution may have contributed to the weakened interaction between D155Y ORF3a and caveolin-1.

**Fig. 7.**
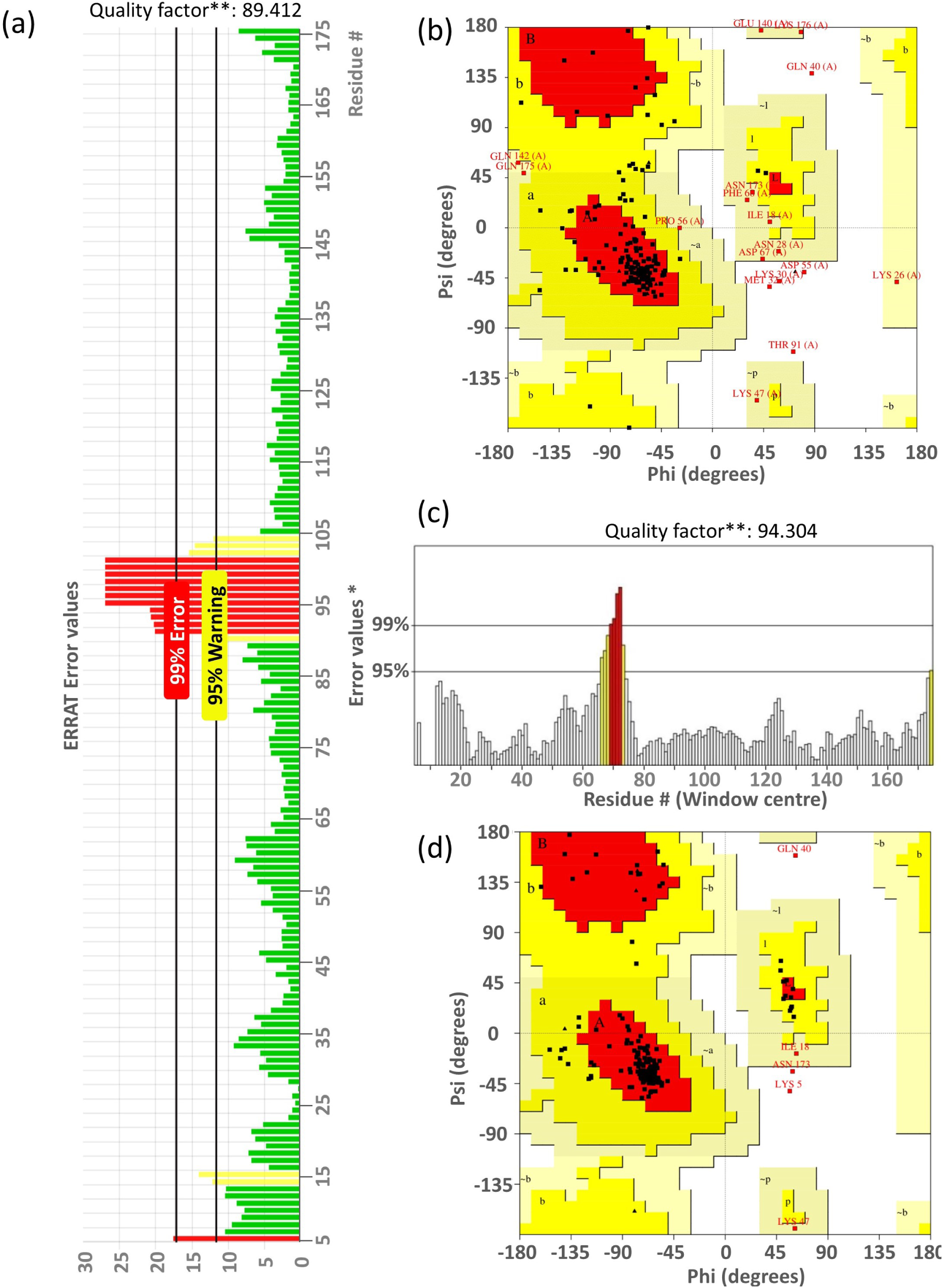
Modelling the human caveolin-1 structure. (a) The ERRAT analysis of the modelled structure of caveolin-1. (b) The distribution of the residues of the modelled structure on the Ramachandran Plot (c) The ERRAT analysis of the simulated structure of caveolin-1. (d) The distribution of the residues of the simulated structure on the Ramachandran Plot. On the error axis, two lines are drawn to indicate the confidence with which it is possible to reject regions that exceed the error value. Overall quality factor 94.304, expressed as the percentage of the protein for which the calculated error value falls below the 95% rejection limit. Good high resolution structures generally produces values around 95% or higher. For lower resolutions (2.5 to 3 Å) the average overall quality factor is around 91%.Stability of the ORF3a-caveolin-1 complex. The time evolution of the RMSD of the ORF3a- caveolin-1 complex with respect to the starting structure. We have considered the data from the last 20ns of our simulation (stable trajectory) and have taken snapshots at intervals of 10ps and done our energy analyses. We have repeated our production runs twice and have considered the average of all the simulation sets. (a) Black: WT-caveolin-1, (b) Red: D155Y-caveolin-1 and (c) Green: S171L-caveolin-1.

**Fig. 8.**
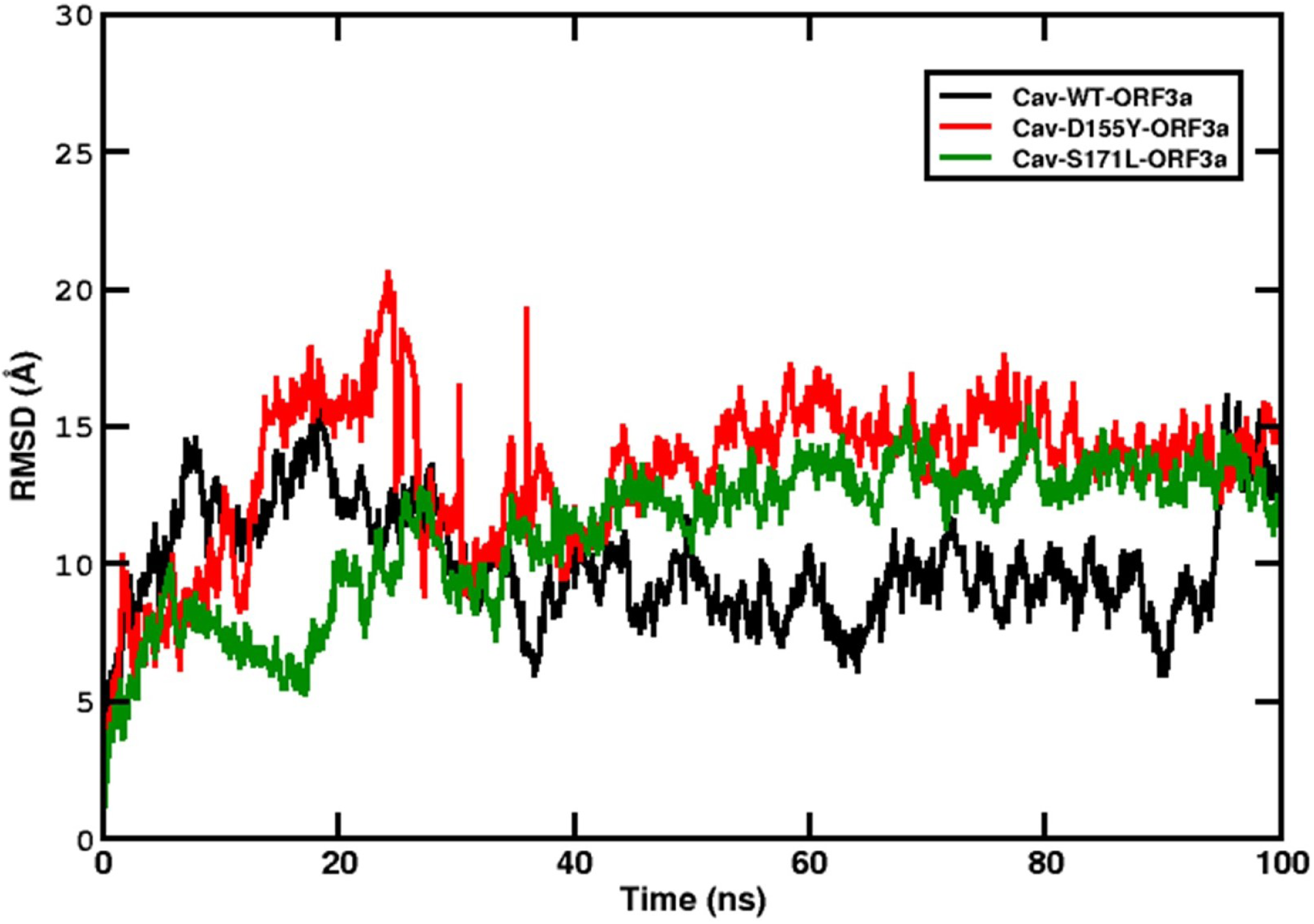
Stability of the ORF3a-caveolin-1 complex. The time evolution of the RMSD of the ORF3a-caveolin-1 complex with respect to the starting structure. We have considered the data from the last 20ns of our simulation (stable trajectory) and have taken snapshots at intervals of 10ps and done our energy analyses. We have repeated our production runs twice (Please see Fig. S6) and have considered the average of all the simulation sets. (a) Black: WT-caveolin-1, (b) Red: D155Y-caveolin-1 and (c) Green: S171L-caveolin-1.

## 4. Discussion

In this study, we provide evidence that substitution D155Y changes the intramolecular hydrogen bond formation, salt bridge formation, and disrupts the interaction between ORF3a and caveolin-1.

We found that several other mutations exist in ORF3a of SARS-CoV-2. The incidence of mutation at position 57 remains high all over the World (as shown in Fig. 1) compared to the other positions (25). But overall percentage of its occurrence was low in India. However, we introduced substitution Q57H in D155Y and another variant S171L, and simulated them for 200 ns. Only the Q57H-D155Y variant was found to be considerably less stable. We also checked the structural stability of W131C, W131R, G172C and G172V, which were found in Domain III and VI, respectively. We simulated the variants W131C, W131R, G172C and G172V for 200 ns. All of these four variants were stable and showed an average RMSD value of 2.5Å with respect to the starting structure as shown in Fig. S4. This indicates that these four mutants are very stable and can have independent existence. Further study is needed to check the effects of these substitutions both *in silico* and *in vitro*. We may hypothesise that the reduced binding affinity of ORF3a to caveolin-1 may be attributed to the structural instability of the D155Y variant. The disrupted interaction may be indicative of improved viral fitness, wherein, the virion particles can continue to build the host intracellular viral load without inducing host cell apoptosis or promoting their egress thus lengthening the asymptomatic phase of the infection. Contrariwise, the ORF3a-caveolin- 1 affinity change can also affect the virion internalisation into host cells, endomembrane sorting and assembly of the viral components.

Direct coupling analysis revealed that eight genes of SARS-CoV-2 genome participate in epistatic interactions at several polymorphic loci. Interestingly, ORF3a showed significant epistatic links with nsp2, nsp6 and nsp12. These intragenetic interactions open up the possibility of potential evolutionary links of the substitutions at D155Y with other positively selected loci in the viral genes, subject to demographic variations (48). Moreover, the Neanderthal-derived COVID-19 risk haplotype is altogether positively selected in some populations and has 30% allele frequency thus introducing an evolutionary landscape to the current COVID-19 pandemic (49). SARS-CoV-2 enters the host cell by both membrane fusion and by clathrin /caveolin-mediated endocytosis after binding to the ACE2 cell-surface receptors in the upper respiratory tract and alveolar epithelial cells (50; 51; 52). It has been hypothesized that internalisation of virion was facilitated by caveolin-1 as several caveolin-1 binding domains (CBD) were found in SARS-CoV protein. However, the role of caveolin-1 was not limited to viral entry. Rather, it was associated with all stages of viral life cycle starting from virus binding to surface receptors, fusion and endo-membrane trafficking of virus in caveosomes, sorting of viral components to endo-membrane surfaces, replication, assembly and to subsequent egress. The host-derived lipid bilayer surrounding the enveloped viral nucleocapsid contains caveolin-1 incorporated during viral fission from the host membrane (53). Thus, change in binding interactions between caveolin-1 and viral proteins containing CBD may provide alternative routes for SARS-CoV-2 pathogenesis.

Previous studies on ORF3a of SARS-CoV have established that multiple domains of ORF3a can interact with different host proteins and modulate various host signalling pathways. Domain I consists of N terminus putative signal peptide (aa 1-15) and it facilitates the translocation of the nascent protein at the ER membrane (19). Domain II (aa 36-40) can interact with TRAF3 and ASC, and activate NLRP3 inflammasome (5). Interestingly, mutation was hardly found in this domain in Indian patients. Domain III has a conserved Cysteine residue at 133^*rd*^ position, which stabilizes ORF3a homodimer and homotetramers for its ion channel activity (46; 54). Further study is needed to understand the importance of mutations at positions 26, 93, 106, 110 and 125, in ion-channel activity of ORF3a. Cytosolic region of domain IV has a conserved motif YDANYFVCW (aa 141-149) that binds with host caveolin-1 (45). Other motifs like *Y XXϕ* and diacidic ExD located in domain V and VI, respectively, participate in intracellular viral protein sorting, trafficking, and its release into culture medium (13). ORF3a has been shown to interact with components of the anti-inflammatory pathway HMOX1, innate immune signalling pathway, TRIM59, glycosylation pathway (ALG5) and nucleus-inner-membrane proteins (SUN2 and ARL6IP6) (55). ORF3a can also regulate Caspase 8-mediated extrinsic apoptotic pathway for its pro-apoptotic activity in HEK293T cells (16). Thus, mutations in the binding regions or in its close proximity may change the interaction between viral ORF3a protein and its binding partner. On the basis of our in silico study, we are currently investigating the effect of D155Y in cell lines. Furthermore, ORF3a elicits significant CD4+ and CD8+ T-cell response. It has been suggested that an optimal vaccine should be inclusive of class I epitopes derived from M, nsp6 and ORF3a (56; 57; 58). Our *in silico* study provides a window to carry out *in vivo* and *in vitro* studies for evaluating the pathogenesis of mutant SARS-CoV-2.

## Supporting information

Revised Supplementary Information

## Author Contribution

PB and SSJ conceptualized the study. SG and PB performed experimental work. SG, PB, DM, KB, SM, SS and SSJ contributed to analysis the results and prepare the figures. SG, DM, KB and SM carried out bioinformatic analysis. SG, DM and KB wrote the initial draft of the manuscript. SSJ, PB and SM edited the manuscript with input from all the authors. All authors agreed to submit the manuscript to the Computational and Structural Biotechnology Journal.

## Conflicts of interest

The authors declare that there are no conflicts of interest.

## Funding information

This study was funded by Technical Research Centre (TRC), Indian Association for the Cultivation of Science (IACS), Kolkata, India.

## Acknowledgement

This work was supported by the Indian Association for the Cultivation of Science (IACS). We thank Google Cloud Research Credits (GCRC) program and the high-performance computing (HPC) support provided by Fluid Numerics. We also thank IACS for fellowship to SG and DM, DST-INSPIRE for fellowship to KB, and SERB-DIA 460 (DIA/2018/000005) award to SSJ.

## Abbreviations

ARL6IP6: ADP Ribosylation Factor Like GTPase 6 interacting protein 6
ASC: Apoptosis associated speck-like protein containing a caspase recruitment domain
BLAST: Basic Local Alignment Search Tool
CD4+: Cluster of Differentiation 4+
CD8+: Cluster of Differentiation 8+
COVID-19: Coronavirus Disease 2019
Cryo-EM: Cryo Electron Microscope
HMOX1: Heme Oxygenase 1
IFN: Interferon
MERS-CoV: Middle East respiratory syndrome coronavirus
MMGBSA: Molecular mechanics with generalized Born and surface area solvation
NCBI: National Centre for Biotechnology Information
NF-*κ*B: Nuclear factor kappa light chain enhancer of activated B cells
NLRP3: Nucleotide-binding oligomerization domain,Leucine rich repeat and Pyrin domain containing
ORF: Open Reading Frame
PDB: Protein Data Bank
PISA: Protein Interfaces Surfaces and Assemblies
PROVEAN: Protein Variation Effect Analyzer
RMSD: Root Mean Square Deviation
SUN2: SUN domain-containing protein 2
TRIM59: Tripartite motif-containing protein 59.

## Footnotes

* PBD files have been provided in the following link - https://github.com/shrimonmuke0202/PDB-File-Protonated-Caveolin-1.git

## Notes

### Competing Interest Statement

The authors have declared no competing interest.

## References

[1] P. V’kovski, A. Kratzel, S. Steiner, H. Stalder, V. Thiel, Coronavirus biology and replication: implications for sars-cov-2, Nature Reviews Microbiology 19 (3) (2021) 155–170.

[2] WHO, Asiaswho indian situation. 2020;10–11 https://www.who.int/india/emergencies/coronavirus-disease-(covid-19)/india-situation-report (2020).

[3] J. Hopkins, Covid-19 resource center https://coronavirus.jhu.edu/map.html (2020).

[4] A. A. T. Naqvi, K. Fatima, T. Mohammad, U. Fatima, I. K. Singh, A. Singh, S. M. Atif, G. Hariprasad, G. M. Hasan, M. I. Hassan, Insights into sars-cov-2 genome, structure, evolution, pathogenesis and therapies: Structural genomics approach, Biochimica et Biophysica Acta (BBA)-Molecular Basis of Disease 1866 (10) (2020) 165878.

[5] K.-L. Siu, K.-S. Yuen, C. Castano-Rodriguez, Z.-W. Ye, M.-L. Yeung, S.-Y. Fung, S. Yuan, C.-P. Chan, K.-Y. Yuen, L. Enjuanes, et al., Severe acute respiratory syndrome coronavirus orf3a protein activates the nlrp3 inflammasome by promoting traf3-dependent ubiquitination of asc, The FASEB Journal 33 (8) (2019) 8865–8877.

[6] A. E. Firth, A putative new sars-cov protein, 3c, encoded in an orf overlapping orf3a, The Journal of general virology 101 (10) (2020) 1085.

[7] Y. Finkel, O. Mizrahi, A. Nachshon, S. Weingarten-Gabbay, D. Morgenstern, Y. Yahalom-Ronen, H. Tamir, H. Achdout, D. Stein, O. Israeli, et al., The coding capacity of sars-cov-2, Nature 589 (7840) (2021) 125– 130.

[8] Y. Konno, I. Kimura, K. Uriu, M. Fukushi, T. Irie, Y. Koyanagi, D. Sauter, R. J. Gifford, S. Nakagawa, K. Sato, et al., Sars-cov-2 orf3b is a potent interferon antagonist whose activity is increased by a naturally occurring elongation variant, Cell reports 32 (12) (2020) 108185.

[9] M. Oostra, C. De Haan, R. De Groot, P. Rottier, Glycosylation of the severe acute respiratory syndrome coronavirus triple-spanning membrane proteins 3a and m, Journal of virology 80 (5) (2006) 2326–2336.

[10] B. Yount, R. S. Roberts, A. C. Sims, D. Deming, M. B. Frieman, J. Sparks, M. R. Denison, N. Davis, R. S. Baric, Severe acute respiratory syndrome coronavirus group-specific open reading frames encode nonessential functions for replication in cell cultures and mice, Journal of virology 79 (23) (2005) 14909–14922.

[11] Y.-J. Tan, The severe acute respiratory syndrome (sars)-coronavirus 3a protein may function as a modulator of the trafficking properties of the spike protein, Virology journal 2 (1) (2005) 1–5.

[12] R. Zeng, R.-F. Yang, M.-D. Shi, M.-R. Jiang, Y.-H. Xie, H.-Q. Ruan, X.-S. Jiang, L. Shi, H. Zhou, L. Zhang, et al., Characterization of the 3a protein of sars-associated coronavirus in infected vero e6 cells and sars patients, Journal of molecular biology 341 (1) (2004) 271–279.

[13] C. Huang, K. Narayanan, N. Ito, C. Peters, S. Makino, Severe acute respiratory syndrome coronavirus 3a protein is released in membranous structures from 3a protein-expressing cells and infected cells, Journal of virology 80 (1) (2006) 210–217.

[14] Y.-J. Tan, P.-Y. Tham, D. Z. Chan, C.-F. Chou, S. Shen, B. C. Fielding, T. H. Tan, S. G. Lim, W. Hong, The severe acute respiratory syndrome coronavirus 3a protein up-regulates expression of fibrinogen in lung epithelial cells, Journal of virology 79 (15) (2005) 10083–10087.

[15] R. Minakshi, K. Padhan, M. Rani, N. Khan, F. Ahmad, S. Jameel, The sars coronavirus 3a protein causes endoplasmic reticulum stress and induces ligand-independent downregulation of the type 1 interferon receptor, PloS one 4 (12) (2009) e8342.

[16] Y. Ren, T. Shu, D. Wu, J. Mu, C. Wang, M. Huang, Y. Han, X.-Y. Zhang, W. Zhou, Y. Qiu, et al., The orf3a protein of sars-cov-2 induces apoptosis in cells, Cellular & molecular immunology 17 (8) (2020) 881– 883.

[17] C. Roy, S. M. Mandal, S. K. Mondal, S. Mukherjee, T. Mapder, W. Ghosh, R. Chakraborty, Trends of mutation accumulation across global sars-cov-2 genomes: Implications for the evolution of the novel coronavirus, Genomics 112 (6) (2020) 5331–5342.

[18] L. Velazquez-Salinas, S. Zarate, S. Eberl, D. P. Gladue, I. Novella, M. V. Borca, Positive selection of orf1ab, orf3a, and orf8 genes drives the early evolutionary trends of sars-cov-2 during the 2020 covid-19 pandemic, Frontiers in Microbiology 11 (2020) 2592.

[19] E. Issa, G. Merhi, B. Panossian, T. Salloum, S. Tokajian, Sars-cov-2 and orf3a: nonsynonymous mutations, functional domains, and viral pathogenesis, Msystems 5 (3) (2020) e00266–20.

[20] A. S. Gonzalez-Reiche, M. M. Hernandez, M. J. Sullivan, B. Ciferri, H. Alshammary, A. Obla, S. Fabre, G. Kleiner, J. Polanco, Z. Khan, et al., Introductions and early spread of sars-cov-2 in the new york city area, Science 369 (6501) (2020) 297–301.

[21] L. Pelkmans, J. Kartenbeck, A. Helenius, Caveolar endocytosis of simian virus 40 reveals a new two-step vesicular-transport pathway to the er, Nature cell biology 3 (5) (2001) 473–483.

[22] G. E. Simmons Jr, H. E. Taylor, J. E. Hildreth, Caveolin-1 suppresses human immunodeficiency virus-1 replication by inhibiting acetylation of nf-κb, Virology 432 (1) (2012) 110–119.

[23] D. Ravid, G. P. Leser, R. A. Lamb, A role for caveolin 1 in assembly and budding of the paramyxovirus parainfluenza virus 5, Journal of virology 84 (19) (2010) 9749–9759.

[24] D. Hailstones, L. S. Sleer, R. G. Parton, K. K. Stanley, Regulation of caveolin and caveolae by cholesterol in mdck cells, Journal of lipid research 39 (2) (1998) 369–379.

[25] F. Wu, S. Zhao, B. Yu, Y.-M. Chen, W. Wang, Z.-G. Song, Y. Hu, Z.W. Tao, J.-H. Tian, Y.-Y. Pei, et al., A new coronavirus associated with human respiratory disease in china, Nature 579 (7798) (2020) 265–269.

[26] Y. Choi, A. P. Chan, Provean web server: a tool to predict the functional effect of amino acid substitutions and indels, Bioinformatics 31 (16) (2015) 2745–2747.

[27] D. M. Kern, B. Sorum, S. S. Mali, C. M. Hoel, S. Sridharan, J. P. Remis, D. B. Toso, A. Kotecha, D. M. Bautista, S. G. Brohawn, Cryoem structure of the sars-cov-2 3a ion channel in lipid nanodiscs, BioRxiv (2020).

[28] E. Krissinel, K. Henrick, Inference of macromolecular assemblies from crystalline state, Journal of molecular biology 372 (3) (2007) 774–797.

[29] N. Guex, M. C. Peitsch, Swiss-model and the swiss-pdb viewer: an environment for comparative protein modeling, electrophoresis 18 (15) (1997) 2714–2723.

[30] J. A. Maier, C. Martinez, K. Kasavajhala, L. Wickstrom, K. E. Hauser, C. Simmerling, ff14sb: improving the accuracy of protein side chain and backbone parameters from ff99sb, Journal of chemical theory and computation 11 (8) (2015) 3696–3713.

[31] D. A. Case, K. Belfon, I. Ben-Shalom, S. R. Brozell, D. Cerutti, T. Cheatham, V. W. D. Cruzeiro, T. Darden, R. E. Duke, G. Giambasu, et al., Amber 2020 (2020).

[32] W. L. Jorgensen, J. Chandrasekhar, J. D. Madura, R. W. Impey, M. L. Klein, Comparison of simple potential functions for simulating liquid water, The Journal of chemical physics 79 (2) (1983) 926–935.

[33] W. Humphrey, A. Dalke, K. Schulten, Vmd: visual molecular dynamics, Journal of molecular graphics 14 (1) (1996) 33–38.

[34] D. R. Roe, T.E. Cheatham III, Ptraj and cpptraj: software for processing and analysis of molecular dynamics trajectory data, Journal of chemical theory and computation 9 (7) (2013) 3084–3095.

[35] B. R. Miller III, T. D. McGee Jr, J. M. Swails, N. Homeyer, H. Gohlke, A. E. Roitberg, Mmpbsa. py: an efficient program for end-state free energy calculations, Journal of chemical theory and computation 8 (9) (2012) 3314–3321.

[36] V. D. Blondel, J.-L. Guillaume, R. Lambiotte, E. Lefebvre, Fast unfolding of communities in large networks, Journal of statistical mechanics: theory and experiment 2008 (10) (2008) P10008.

[37] J. Yang, R. Yan, A. Roy, D. Xu, J. Poisson, Y. Zhang, The i-tasser suite: protein structure and function prediction, Nature methods 12 (1) (2015) 7–8.

[38] J. Yang, Y. Zhang, I-tasser server: new development for protein structure and function predictions, Nucleic acids research 43 (W1) (2015) W174–W181.

[39] A. Roy, A. Kucukural, Y. Zhang, I-tasser: a unified platform for automated protein structure and function prediction, Nature protocols 5 (4) (2010) 725–738.

[40] G. N. Ramachandran, C. Ramakrishnan, V. Sasisekharan, Stereochemistry of polypeptide chain configurations, J Mol Biol 7 (1963) 95–99.

[41] C. Colovos, T. O. Yeates, Verification of protein structures: patterns of nonbonded atomic interactions, Protein science 2 (9) (1993) 1511–1519.

[42] E. F. Pettersen, T. D. Goddard, C. C. Huang, G. S. Couch, D. M. Greenblatt, E. C. Meng, T. E. Ferrin, Ucsf chimera—a visualization system for exploratory research and analysis, Journal of computational chemistry 25 (13) (2004) 1605–1612.

[43] C. Dominguez, R. Boelens, A. M. Bonvin, Haddock: a proteinprotein docking approach based on biochemical or biophysical information, Journal of the American Chemical Society 125 (7) (2003) 1731–1737.

[44] Q.-C. Cai, Q.-W. Jiang, G.-M. Zhao, Q. Guo, G.-W. Cao, T. Chen, Putative caveolin-binding sites in sars-cov proteins, Acta pharmacologica Sinica 24 (10) (2003) 1051–1059.

[45] K. Padhan, C. Tanwar, A. Hussain, P. Y. Hui, M. Y. Lee, C. Y. Cheung, J. S. M. Peiris, S. Jameel, Severe acute respiratory syndrome coronavirus orf3a protein interacts with caveolin, Journal of General Virology 88 (11) (2007) 3067–3077.

[46] W. Lu, B.-J. Zheng, K. Xu, W. Schwarz, L. Du, C. K. Wong, J. Chen, S. Duan, V. Deubel, B. Sun, Severe acute respiratory syndrome-associated coronavirus 3a protein forms an ion channel and modulates virus release, Proceedings of the National Academy of Sciences 103 (33) (2006) 12540–12545.

[47] N. S. Pagadala, K. Syed, J. Tuszynski, Software for molecular docking: a review, Biophysical Reviews 9 (2) (2017) 91–102. doi:10.1007/s12551-016-0247-1. URL https://doi.org/10.1007/s12551-016-0247-1

[48] H.-L. Zeng, V. Dichio, E. R. Horta, K. Thorell, E. Aurell, Global analysis of more than 50,000 sars-cov-2 genomes reveals epistasis between eight viral genes, Proceedings of the National Academy of Sciences 117 (49) (2020) 31519–31526.

[49] H. Zeberg, S. Pääbo, The major genetic risk factor for severe covid-19 is inherited from neanderthals, Nature 587 (7835) (2020) 610–612.

[50] Y. Inoue, N. Tanaka, Y. Tanaka, S. Inoue, K. Morita, M. Zhuang, T. Hattori, K. Sugamura, Clathrin-dependent entry of severe acute respiratory syndrome coronavirus into target cells expressing ace2 with the cytoplasmic tail deleted, Journal of virology 81 (16) (2007) 8722–8729.

[51] R. Wölfel, V. M. Corman, W. Guggemos, M. Seilmaier, S. Zange, M. A. Müller, D. Niemeyer, T. C. Jones, P. Vollmar, C. Rothe, et al., Virological assessment of hospitalized patients with covid-2019, Nature 581 (7809) (2020) 465–469.

[52] K. Fecchi, S. Anticoli, D. Peruzzu, E. Iessi, M. C. Gagliardi, P. Matarrese, A. Ruggieri, Coronavirus interplay with lipid rafts and autophagy unveils promising therapeutic targets, Frontiers in Microbiology 11 (2020) 1821.

[53] Y. Xing, Z. Wen, W. Gao, Z. Lin, J. Zhong, Y. Jiu, Multifaceted functions of host cell caveolae/caveolin-1 in virus infections, Viruses 12 (5) (2020) 487.

[54] V. Marquez-Miranda, M. Rojas, Y. Duarte, I. Diaz-Franulic, M. Holmgren, R. Cachau, F. D. Gonzalez-Nilo, Analysis of sars-cov-2 orf3a structure reveals chloride binding sites, bioRxiv (2020).

[55] D. E. Gordon, G. M. Jang, M. Bouhaddou, J. Xu, K. Obernier, K. M. White, M. J. O’Meara, V. V. Rezelj, J. Z. Guo, D. L. Swaney, et al., A sars-cov-2 protein interaction map reveals targets for drug repurposing, Nature 583 (7816) (2020) 459–468.

[56] J. Mateus, et al., Selective cross-reactive sars-cov-2 t cell epitopes in unexposed humans [published online august 4, 2020], Science 10.

[57] A. Grifoni, D. Weiskopf, S. I. Ramirez, J. Mateus, J. M. Dan, C. R. Moderbacher, S. A. Rawlings, A. Sutherland, L. Premkumar, R. S. Jadi, et al., Targets of t cell responses to sars-cov-2 coronavirus in humans with covid-19 disease and unexposed individuals, Cell 181 (7) (2020) 1489–1501.

[58] N. Le Bert, A. T. Tan, K. Kunasegaran, C. Y. Tham, M. Hafezi, A. Chia, M. H. Y. Chng, M. Lin, N. Tan, M. Linster, et al., Sars-cov-2-specific t cell immunity in cases of covid-19 and sars, and uninfected controls, Nature 584 (7821) (2020) 457–462.

[59] M. Bastian, S. Heymann, M. Jacomy, Gephi: an open source software for exploring and manipulating networks, in: Third international AAAI conference on weblogs and social media, 2009.

