## Supplementary material for "D155Y Substitution of SARS-CoV-2 ORF3a Weakens Binding with Caveolin-1": Revised Supplementary Information

Table S1: Number of samples showing mutations in ORF3a protein

| Beginning of the table | | | | | | | |
|---|---|---|---|---|---|---|---|
| position # | # of Mutations (world) | # of Mutations (India) | % of occurrence (India) | position # | # of Mutations (world) | # of Mutations (India) | % of occurrence (India) |
| 1 | 0 | 0 | - | 35 | 327 | 1 | 0.306 |
| 2 | 126 | 0 | 0 | 36 | 166 | 0 | 0 |
| 3 | 23 | 0 | 0 | 37 | 319 | 0 | 0 |
| 4 | 2 | 0 | 0 | 38 | 1979 | 0 | 0 |
| 5 | 40 | 3 | 7.5 | 39 | 512 | 0 | 0 |
| 6 | 68 | 1 | 1.471 | 40 | 636 | 1 | 0.157 |
| 7 | 120 | 0 | 0 | 41 | 735 | 6 | 0.816 |
| 8 | 13 | 0 | 0 | 42 | 37308 | 4 | 0.011 |
| 9 | 87 | 0 | 0 | 43 | 89 | 0 | 0 |
| 10 | 524 | 0 | 0 | 44 | 583 | 0 | 0 |
| 11 | 61 | 0 | 0 | 45 | 358 | 1 | 0.279 |
| 12 | 331 | 0 | 0 | 46 | 299 | 0 | 0 |
| 13 | 516 | 2 | 0.388 | 47 | 275 | 0 | 0 |
| 14 | 245 | 0 | 0 | 48 | 170 | 0 | 0 |
| 15 | 6746 | 6 | 0.0889 | 49 | 346 | 0 | 0 |
| 16 | 467 | 3 | 0.6423 | 50 | 118 | 1 | 0.847 |
| 17 | 44 | 0 | 0 | 51 | 186 | 0 | 0 |
| 18 | 566 | 5 | 0.883 | 52 | 232 | 10 | 4.310 |
| 19 | 189 | 1 | 0.529 | 53 | 582 | 2 | 0.344 |
| 20 | 189 | 0 | 0 | 54 | 3442 | 2 | 0.058 |
| 21 | 903 | 2 | 0.221 | 55 | 628 | 4 | 0.637 |
| 22 | 456 | 3 | 0.658 | 56 | 52 | 0 | 0 |
| 23 | 1464 | 2 | 0.137 | 57 | 155611 | 471 | 0.303 |
| 24 | 217 | 0 | 0 | 58 | 1111 | 1 | 0.090 |
| 25 | 217 | 0 | 0 | 59 | 530 | 2 | 0.377 |
| 26 | 28662 | 396 | 1.382 | 60 | 866 | 0 | 0 |
| 27 | 2643 | 0 | 0 | 61 | 63 | 0 | 0 |
| 28 | 123 | 0 | 0 | 62 | 747 | 0 | 0 |
| 29 | 207 | 0 | 0 | 63 | 313 | 0 | 0 |
| 30 | 177 | 0 | 0 | 64 | 240 | 0 | 0 |
| 31 | 217 | 0 | 0 | 65 | 346 | 5 | 1.445 |
| 32 | 468 | 0 | 0 | 66 | 125 | 1 | 0.8 |
| 33 | 3529 | 0 | 0 | 67 | 541 | 0 | 0 |
| 34 | 147 | 0 | 0 | 68 | 1050 | 0 | 0 |

| Continuation of Table 1 | | | | | | | |
|---|---|---|---|---|---|---|---|
| position # | # of Mutations (world) | # of Mutations (India) | % of occurrence (India) | position # | # of Mutations (world) | # of Mutations (India) | % of occurrence (India) |
| 69 | 1050 | 0 | 0 | 109 | 57 | 1 | 1.754 |
| 70 | 35 | 0 | 0 | 110 | 1243 | 17 | 1.368 |
| 71 | 69 | 0 | 0 | 111 | 10 | 0 | 0 |
| 72 | 384 | 1 | 0.260 | 112 | 817 | 1 | 0.122 |
| 73 | 549 | 1 | 0.182 | 113 | 51 | 0 | 0 |
| 74 | 1459 | 0 | 0 | 114 | 30 | 0 | 0 |
| 75 | 183 | 0 | 0 | 115 | 18 | 2 | 11.11 |
| 76 | 58 | 0 | 0 | 116 | 58 | 0 | 0 |
| 77 | 429 | 5 | 1.166 | 117 | 24 | 0 | 0 |
| 78 | 1099 | 0 | 0 | 118 | 91 | 0 | 0 |
| 79 | 10 | 0 | 0 | 119 | 32 | 0 | 0 |
| 80 | 40 | 3 | 7.5 | 120 | 36 | 0 | 0 |
| 81 | 102 | 0 | 0 | 121 | 20 | 0 | 0 |
| 82 | 16 | 0 | 0 | 122 | 101 | 0 | 0 |
| 83 | 185 | 0 | 0 | 123 | 117 | 0 | 0 |
| 84 | 36 | 0 | 0 | 124 | 66 | 0 | 0 |
| 85 | 719 | 2 | 0.278 | 125 | 455 | 13 | 2.857 |
| 86 | 456 | 1 | 0.219 | 126 | 245 | 0 | 0 |
| 87 | 29 | 0 | 0 | 127 | 309 | 0 | 0 |
| 88 | 420 | 2 | 0.476 | 128 | 463 | 4 | 0.864 |
| 89 | 1015 | 1 | 0.098 | 129 | 365 | 0 | 0 |
| 90 | 1123 | 0 | 0 | 130 | 25 | 0 | 0 |
| 91 | 11 | 0 | 0 | 131 | 6122 | 6 | 0.098 |
| 92 | 663 | 0 | 0 | 132 | 13 | 0 | 0 |
| 93 | 286 | 16 | 5.594 | 133 | 14 | 0 | 0 |
| 94 | 413 | 0 | 0 | 134 | 1513 | 4 | 0.264 |
| 95 | 265 | 2 | 0.755 | 135 | 6 | 0 | 0 |
| 96 | 116 | 0 | 0 | 136 | 40 | 0 | 0 |
| 97 | 239 | 1 | 0.418 | 137 | 25 | 1 | 4 |
| 98 | 186 | 0 | 0 | 138 | 3 | 0 | 0 |
| 99 | 1903 | 0 | 0 | 139 | 10 | 1 | 10 |
| 100 | 3228 | 2 | 0.062 | 140 | 521 | 1 | 0.192 |
| 101 | 253 | 1 | 0.395 | 141 | 5 | 0 | 0 |
| 102 | 81 | 0 | 0 | 142 | 75 | 0 | 0 |
| 103 | 427 | 1 | 0.234 | 143 | 381 | 1 | 0.262 |
| 104 | 9158 | 3 | 0.033 | 144 | 34 | 0 | 0 |
| 105 | 44 | 1 | 2.273 | 145 | 44 | 0 | 0 |
| 106 | 693 | 15 | 2.345 | 146 | 3 | 1 | 33.33 |
| 107 | 312 | 0 | 0 | 147 | 197 | 0 | 0 |
| 108 | 1350 | 2 | 0.148 | 148 | 27 | 2 | 7.407 |

| Continuation of Table 1 | | | | | | | |
|---|---|---|---|---|---|---|---|
| position # | # of Mutations (world) | # of Mutations (India) | % of occurrence (India) | position # | # of Mutations (world) | # of Mutations (India) | % of occurrence (India) |
| 149 | 154 | 1 | 0.650 | 189 | 109 | 2 | 1.835 |
| 150 | 55 | 0 | 0 | 190 | 33 | 0 | 0 |
| 151 | 3111 | 3 | 0.096 | 191 | 78 | 0 | 0 |
| 152 | 58 | 1 | 1.724 | 192 | 39 | 0 | 0 |
| 153 | 96 | 0 | 0 | 193 | 461 | 0 | 0 |
| 154 | 82 | 0 | 0 | 194 | 21 | 0 | 0 |
| 155 | 2261 | 33 | 1.460 | 195 | 127 | 0 | 0 |
| 156 | 23 | 0 | 0 | 196 | 857 | 0 | 0 |
| 157 | 1 | 0 | 0 | 197 | 51 | 0 | 0 |
| 158 | 134 | 0 | 0 | 198 | 136 | 0 | 0 |
| 159 | 209 | 0 | 0 | 199 | 28 | 0 | 0 |
| 160 | 6 | 0 | 0 | 200 | 34 | 0 | 0 |
| 161 | 6 | 0 | 0 | 201 | 68 | 0 | 0 |
| 162 | 113 | 2 | 1.770 | 202 | 829 | 0 | 0 |
| 163 | 162 | 0 | 0 | 203 | 5 | 0 | 0 |
| 164 | 7 | 0 | 0 | 204 | 30 | 0 | 0 |
| 165 | 248 | 0 | 0 | 205 | 31 | 0 | 0 |
| 166 | 974 | 3 | 0.308 | 206 | 27 | 0 | 0 |
| 167 | 8 | 1 | 12.5 | 207 | 188 | 0 | 0 |
| 168 | 35 | 0 | 0 | 208 | 15 | 0 | 0 |
| 169 | 33 | 0 | 0 | 209 | 55 | 0 | 0 |
| 170 | 18 | 0 | 0 | 210 | 484 | 2 | 0.413 |
| 171 | 3147 | 12 | 0.381 | 211 | 51 | 0 | 0 |
| 172 | 51165 | 5 | 0.010 | 212 | 0 | 0 | - |
| 173 | 331 | 6 | 1.813 | 213 | 346 | 0 | 0 |
| 174 | 2482 | 3 | 0.121 | 214 | 1 | 0 | 0 |
| 175 | 1519 | 2 | 0.132 | 215 | 6 | 0 | 0 |
| 176 | 407 | 0 | 0 | 216 | 67 | 0 | 0 |
| 177 | 222 | 1 | 0.450 | 217 | 351 | 0 | 0 |
| 178 | 26 | 0 | 0 | 218 | 90 | 0 | 0 |
| 179 | 41 | 0 | 0 | 219 | 290 | 0 | 0 |
| 180 | 2355 | 1 | 0.042 | 220 | 123 | 1 | 0.813 |
| 181 | 114 | 0 | 0 | 221 | 106 | 0 | 0 |
| 182 | 158 | 10 | 6.329 | 222 | 114 | 0 | 0 |
| 183 | 551 | 0 | 0 | 223 | 5066 | 37 | 0.730 |
| 184 | 66 | 0 | 0 | 224 | 2346 | 7 | 0.298 |
| 185 | 708 | 0 | 0 | 225 | 1107 | 0 | 0 |
| 186 | 153 | 0 | - | 226 | 661 | 1 | 0.151 |
| 187 | 49 | 0 | 0 | 227 | 90 | 1 | 1.111 |
| 188 | 2581 | 0 | 0 | 228 | 38 | 0 | 0 |

| Continuation of Table 1 | | | | | | | |
|---|---|---|---|---|---|---|---|
| position # | # of Mutations (world) | # of Mutations (India) | % of occurrence (India) | position # | # of Mutations (world) | # of Mutations (India) | % of occurrence (India) |
| 229 | 368 | 0 | 0 | 255 | 86 | 0 | 0 |
| 230 | 75 | 0 | 0 | 256 | 310 | 0 | 0 |
| 231 | 22 | 0 | 0 | 257 | 725 | 1 | 0.138 |
| 232 | 60 | 0 | 0 | 258 | 103 | 0 | 0 |
| 233 | 4 | 0 | 0 | 259 | 240 | 0 | 0 |
| 234 | 19 | 0 | 0 | 260 | 330 | 0 | 0 |
| 235 | 63 | 0 | 0 | 261 | 92 | 0 | 0 |
| 236 | 828 | 0 | 0 | 262 | 448 | 2 | 0.446 |
| 237 | 87 | 0 | 0 | 263 | 50 | 3 | 6 |
| 238 | 260 | 3 | 1.154 | 264 | 99 | 0 | 0 |
| 239 | 3536 | 2 | 0.056 | 265 | 62 | 0 | 0 |
| 240 | 7589 | 3 | 0.040 | 266 | 63 | 1 | 1.587 |
| 241 | 312 | 1 | 1.111 | 267 | 784 | 0 | 0 |
| 242 | 60 | 0 | 0 | 268 | 362 | 1 | 0.276 |
| 243 | 15 | 1 | 6.667 | 269 | 455 | 0 | 0 |
| 244 | 32 | 0 | 0 | 270 | 457 | 0 | 0 |
| 245 | 66 | 1 | 1.515 | 271 | 64 | 0 | 0 |
| 246 | 49 | 0 | 0 | 272 | 112 | 0 | 0 |
| 247 | 70 | 0 | 0 | 273 | 102 | 0 | 0 |
| 248 | 7 | 0 | 0 | 274 | 18 | 1 | 5.556 |
| 249 | 3 | 0 | 0 | 275 | 213 | 3 | 1.408 |
| 250 | 6 | 1 | 16.667 | - | - | - | - |
| 251 | 674 | 0 | 0 | - | - | - | - |
| 252 | 147 | 0 | 0 | - | - | - | - |
| 253 | 15954 | 0 | 0 | - | - | - | - |
| 254 | 748 | 0 | 0 | - | - | - | - |
| End of Table | | | | | | | |

Table S2: The list of residues forming hydrogen bonds showing maximum occupancy in the three systems are shown. Main and side indicate main chain and side chain hydrogen bonds.

| Residue1 | Residue2 | Occupancy |
|---|---|---|
| TYR156'-Side | LYS192'-Main | 71.93%±0.0019 |
| ARG134-Side | ASP155-Side | 70.1%±0.0084 |
| SER205'-Side | ASN144'-Main | 67.99%±0.0022 |
| LEU203-Main | ASP210-Main | 68.28%±0.0013 |
| SER205-Side | ASN144-Main | 67.86%±0.0020 |
| LEU203'-Main | ASP210'-Main | 67.97%±0.0010 |
| TYR154-Side | GLU191-Side | 61.90%±0.0005 |
| THR89'-Side | LEU85'-Main | 61.75%±0.0011 |
| THR89-Side | LEU85-Main | 60.97%±0.0021 |
| THR175'-Side | ASP173'-Side | 61.7%±0.0036 |

(a) WT

| Residue1 | Residue2 | Occupancy |
|---|---|---|
| TYR212'-Side | THR164'-Main | 68.44%±0.0129 |
| SER205'-Side | ASN144'-Main | 69.27%±0.0022 |
| SER205-Side | ASN144-Main | 68.99%±0.0014 |
| TYR233'-Side | LYS198'-Main | 66.44%±0.0006 |
| LEU203-Main | ASP210-Main | 66.20%±0.0035 |
| TYR233-Side | LYS198-Main | 66.44%±0.0035 |
| LEU203'-Main | ASP210'-Main | 66.86%±0.0013 |
| THR89'-Side | LEU85'-Main | 63.12%±0.0021 |
| TYR233-Main | ILE167-Main | 59.67%±0.0077 |
| THR89-Side | LEU85-Main | 61.11%±0.0005 |

(b) D155Y

| Residue1 | Residue2 | Occupancy |
|---|---|---|
| LEU203-Main | ASP210-Main | 64.48%±0.0074 |
| LEU203'-Main | ASP210'-Main | 63.23%±0.0018 |
| THR89-Side | LEU85'-Main | 63.37%±0.0068 |
| TYR233'-Side | LYS198'-Main | 61.92%±0.0121 |
| TYR91-Side | PRO42-Main | 55.81%±0.0182 |
| SER58'-Side | ALA54'-Main | 62.31%±0.0085 |
| THR89-Side | LEU85-Main | 59.91%±0.0088 |
| TYR154'-Side | GLU191'-Side | 60.33%±0.0072 |
| TYR233-Side | LYS198-Main | 59.93%±0.0011 |
| SER58-Side | ALA54-Main | 61.28%±0.0027 |

(c) S171L

Table S3: List of salt bridge interactions in the three systems

| Salt Bridge | WT | D155Y | S171L | Salt Bridge | WT | D155Y | S171L |
|---|---|---|---|---|---|---|---|
| ASP142-LYS61’ | ✓ | ✓ | ✓ | ASP199-LYS198 | ✗ | ✓ | ✓ |
| ASP142-LYS75 | ✓ | ✓ | ✓ | ASP199’-LYS198’ | ✓ | ✓ | ✓ |
| ASP142-ARG116 | ✓ | ✗ | ✗ | ASP210-LYS235 | ✓ | ✓ | ✓ |
| ASP142-ARG122 | ✗ | ✓ | ✓ | ASP210’-LYS235’ | ✓ | ✓ | ✓ |
| ASP142-ARG126 | ✗ | ✓ | ✓ | ASP238-LYS235 | ✓ | ✓ | ✓ |
| ASP142’-LYS61 | ✓ | ✓ | ✓ | ASP238’-LYS235’ | ✓ | ✓ | ✓ |
| ASP142’-ARG122’ | ✓ | ✗ | ✓ | GLU181-LYS192 | ✓ | ✓ | ✓ |
| ASP142’-LYS61’ | ✓ | ✗ | ✗ | GLU181-LYS67 | ✗ | ✗ | ✓ |
| ASP142’-LYS75’ | ✓ | ✓ | ✓ | GLU181-ARG67 | ✗ | ✗ | ✓ |
| ASP142’-ARG116’ | ✓ | ✗ | ✗ | GLU181’-LYS192’ | ✓ | ✓ | ✓ |
| ASP142’-LYS75 | ✓ | ✗ | ✓ | GLU181’-LYS266 | ✓ | ✗ | ✗ |
| ASP142’-ARG126’ | ✗ | ✓ | ✓ | GLU191-ARG68 | ✓ | ✓ | ✓ |
| ASP155-ARG134 | ✓ | ✗ | ✓ | GLU191-LYS192 | ✗ | ✗ | ✓ |
| ASP155’-ARG134’ | ✓ | ✗ | ✓ | GLU191’-LYS192’ | ✓ | ✗ | ✗ |
| ASP181’-LYS192’ | ✗ | ✓ | ✗ | GLU191’-ARG68’ | ✓ | ✓ | ✓ |
| ASP183-LYS192 | ✓ | ✓ | ✓ | GLU194-LYS192 | ✓ | ✓ | ✓ |
| ASP183-LYS67 | ✓ | ✓ | ✓ | GLU194-LYS198 | ✗ | ✗ | ✓ |
| ASP183-ARG68 | ✓ | ✗ | ✓ | GLU194’-LYS192’ | ✓ | ✓ | ✓ |
| ASP183’-LYS67’ | ✓ | ✗ | ✗ | GLU226-LYS235’ | ✓ | ✗ | ✗ |
| ASP183’-LYS192’ | ✗ | ✗ | ✓ | GLU226’-LYS23 | ✓ | ✗ | ✗ |
| ASP199-LYS1 | ✓ | ✗ | ✗ | GLU226’-LYS235 | ✗ | ✓ | ✓ |

Table S4: List of residues constituting each cluster for WT and the two mutants. The functional domain in which some of the constituent residues fall are indicated within parentheses.

| Beginning of the table | | |
|---|---|---|
| WT | D155Y | S171L |
| 0: ALA51 ALA54 ALA59 GLN57 GLY49 GLY44 ILE47 LEU52 LEU53 LEU41 LEU46 PHE56 PHE43 PRO42 SER40 SER58 SER60 TRP45 VAL50 VAL55 VAL48 | 0: ALA51 ALA54 ALA59 GLN57 GLY49 GLY44 ILE62 ILE47 LEU52 LEU53 LEU41 LEU46 LYS61 PHE56 PHE43 PRO42 SER40 SER58 SER60 TRP45 VAL50 VAL55 VAL48 | 0: ALA51 ALA54 ALA59 ASN82 CYS81 GLN57 GLY49 GLY76 GLY44 HIE78 HIE93 ILE47 LEU52 LEU53 LEU41 LEU83 LEU84 LEU85 LEU86 LEU94 LEU95 LEU46 LYS75 PHE56 PHE43 PHE79 PHE87 PRO42 SER40 SER58 SER60 SER92 THR89 TRP45 TYR91 VAL50 VAL55 VAL77 VAL80 VAL88 VAL90 VAL48 |
| 1: ALA51’ ALA54’ ALA59’ ASN82’ CYS81’ GLN57’ GLY44’ GLY49’ GLY76’ HIE78’ HIE93’ ILE47’ LEU41’ LEU46’ LEU52’ LEU53’ LEU83’ LEU84’ LEU85’ LEU86’ LEU94’ LEU95’ LYS75’ PHE43’ PHE56’ PHE79’ PHE87’ PRO42’ SER40’ SER58’ SER60’ SER92’ THR89’ TRP45’ TYR91’ VAL48’ VAL50’ VAL55’ VAL77’ VAL80’ VAL88’ VAL90’ (Domain 3) | 1: ALA98 ALA99 ASN82 CYS81 GLY76 GLY100 HIE78 HIE93 LEU83 LEU84 LEU85 LEU86 LEU94 LEU95 LEU96 LYS75 PHE79 PHE87 SER92 THR89 TYR91 VAL77 VAL80 VAL88 VAL90 VAL97 (Domain 3) | 1: ALA51’ ALA52’ ALA59’ GLN57’ GLY44’ GLY49’ ILE47’ ILE62’ LEU41’ LEU46’ LEU52’ LEU53’ LYS61’ PHE43’ PHE56’ PRO42’ SER40’ SER58’ SER60’ TRP45’ VAL48’ VAL50’ VAL55’ VAL50’ VAL55’ |
| 2: ASN82 CYS81 GLY76 HIE78 HIE93 LEU83 LEU84 LEU85 LEU86 LEU94 LEU95 LYS75 PHE79 PHE87 SER92 THR89 TYR91 VAL77 VAL80 VAL88 VAL90 | 2: ALA51’ ALA54’ ALA59’ GLN57’ GLY44’ GLY49’ ILE47’ ILE62’ ILE63’ LEU41’ LEU44’ LEU52 LEU53’ LYS61’ PHE43’ PHE56’ PRO42’ SER40’ SER58’ SER60’ THR64’ TRP45’ VAL48’ VAL50’ VAL55’ | 2: ALA62’ ARG68’ ASN82’ CYS81’ GLN70’ GLY76’ HIE78’ HIE93’ ILE63’ LEU65’ LEU71’ LEU83 LEU84’ LEU85’ LEU86’ LEU94’ LEU95’ LYS66’ LYS67’ LYS75’ PHE79’ PHE87’ SER74’ SER92’ THR64’ THR89’ TRP69’ TYR91’ VAL77’ VAL80’ VAL88’ VAL90’ |

| Continuation of Table S4 | | |
|---|---|---|
| WT | D155Y | S171L |
| 3: ALA88 ALA99 ALA103 ALA110 ARG122 ASN119 GLN116 GLU102 GLY100 ILE118 ILE123 ILE124 LEU96 LEU101 LEU106 LEU108 LEU111 LEU115 MET125 PHE105 PHE114 PHE120 PRO104 SER117 TYR107 TYR109 TYR113 VAL97 VAL112 VAL121 (Domain 3) | 3: ALA98’ ALA99’ ALA103’ ALA110’ ARG122’ ASN119’ GLN116’ GLU102’ GLY100’ ILE118’ ILE123’ ILE124’ LEU96’ LEU101’ LEU106 LEU108’ LEU111’ LEU115’ MET125’ PHE105’ PHE114’ PHE120’ PRO104’ SER117 TYR107’ TYR109’ TYR113’ VAL97’ VAL112’ VAL121’ (Domain 3) | 3: ALA98’ ALA99’ ALA103’ ALA110’ ARG122’ ASN119’ GLN116’ GLU102’ GLY100 ILE118’ ILE123’ ILE124’ LEU96’ LEU101’ LEU106’ LEU108’ LEU109’ LEU115 MET125’ PHE105’ PHE114’ PHE120’ PRO104’ SER117’ TYR107’ TYR109’ TYR113’ VAL97’ VAL112’ VAL121’ (Domain 3) |
| 4: ALA88’ ALA99’ ALA103’ ALA110’ GLN106’ GLU102’ GLY100’ LEU96’ LEU101’ LEU106’ LEU108’ LEU111’ LEU115’ PHE105’ PHE114’ PRO104’ SER117’ TYR107’ TYR109’ TYR113’ VAL97’ VAL112’ (Domain 3) | 4: ALA72’ ARG68’ ASN82’ CYS81’ GLN70’ GLY76’ HIE78’ HIE93’ LEU65’ LEU71’ LEU73’ LEU83’ LEU84’ LEU85’ LEU86’ LEU94’ LEU95’ LYS66’ LYS67’ LYS75’ PHE79’ PHE87’ SER74’ SER92’ THR89’ TRP69’ TYR91’ VAL77’ VAL80’ VAL88’ VAL90’ (Domain 3) | 4: ALA98 ALA99 ALA103 ALA110 ASN119 GLN116 GLU102 GLY100 ILE118 LEU96 LEU101 LEU101’ LEU108 LEU111 LEU115 PHE105 PHE114 PHE120 PRO104 SER117 TYR107 TYR109 TYR113 VAL97 VAL112 VAL121 (Domain 3) |
| 5: ALA72 ARG68 GLN70 ILE62 ILE63 LEU65 LEU71 LEU73 LYS61 LYS66 LYS67 SER74 THR64 TRP69 | 5: ALA103 ALA11 ARG122 ASN119 GLN116 GLU102 ILE118 ILE123 ILE124 LEU101 LEU106 LEU108 LEU111 LEU115 MET125 PHE105 PHE114 PHE120 PRO104 SER117 TYR107 TYR109 TYR113 VAL112 VAL121 (Domain 3) | 5: ALA72 ARG68 GLN70 ILE62 ILE63 LEU65 LEU71 LEU73 LYS61 LYS66 LYS67 SER74 THR64 TRP69 |
| 6: ARG122’ ARG126’ ASN119’ ASN119’ ASP199’ ASP210’ ASP238’ CYS130’ CYS133’ CYS200’ HIE204’ ILE118’ ILE123’ ILE124’ ILE236’ LYS132’ LYS198’ LYS235’ MET125’ PHE120’ PHE207’ SER205’ SER209’ THR208’ TRP127’ TRP131’ TYR206’ TYR211’ TYR212’ VAL121’ VAL200’ VAL202’ VAL237’ (Domain 3) | 6: ALA72 ARG68 GLN70 ILE63 LEU65 LEU71 LEU73 LYS66 LYS67 SER74 THR64 TRP69 | 6: ALA143’ ARG126’ ARG134’ ASN137’ ASN144’ ASN152’ ASP142’ ASP155’ ASP199’ ASP210’ ASP238’ CYS130’ CYS133’ CYS148’ CYS153’ CYS157’ CYS200’ HIE150’ HIE204’ ILE158’ ILE236’ LEU127’ LEU129’ LEU139’ LEU140’ LEU147’ LEU203’ LYS132’ LYS136’ LYS198’ LYS235’ PHE146’ PHE207’ PRO138’ SER135’ SER205’ |

| Continuation of Table S4 | | |
|---|---|---|
| WT | D155Y | S171L |
| | | SER209’ THR151’ THR208 TRP128’ TRP131’ TRP149’ TYR141’ TYR145’ TYR154 TYR156’ TYR206’ TYR211’ TYR212’ VAL201’ VAL202’ VAL237’ (Domain 4) |
| 7: ALA72’ ARG68’ GLN70’ ILE62’ ILE63’ LEU65’ LEU71’ LEU73’ LYS61’ LEU71’ LEU73’ LYS61’ LYS66’ LYS67’ SER74’ THR64’ TRP69’ | 7: ALA143’ ARG126’ ARG134’ ASN137’ ASN144’ ASN152’ ASN161’ ASP142’ ASP199’ ASP210’ ASP238’ CYS130’ CYS133’ CYS148’ CYS153’ CYS157’ CYS200’ HIE150’ HIE204’ ILE158’ ILE236’ LEU127’ LEU129’ LEU139’ LEU140’ LEU147’ LEU203’ LYS142’ LYS146’ LYS198’ LYS235’ PHE146’ PHE207’ PRO138’ PRO159’ SER135’ SER162’ SER165’ SER166’ SER205’ SER209’ THR151’ THR164’ THR208’ TRP128’ TRP131’ TRP149’ TYR141’ TYR145’ TYR154’TYR155’ TYR156’ TYR160’ TYR206’ TYR211’ TYR212’ VAL163’ VAL210’ VAL202’ VAL237’ (Domains 3, 4, 5) | 7: ASN161’ ASN234’ ASP222’ GLN213’ GLN218’ GLU226’ GLY224’ HIE227’ ILE167’ ILE169’ ILE232’ LEU214’ LEU219’ PHE230’ PHE231’ PRO159’ SER162’ SER165’ SER166’ SER216’ SER220’ THR164’ THR170’ THR217’ THR221’ THR223’ THR229’ TYR160’ TYR215’ TYR133’ VAL163’ VAL168’ VAL225’ VAL228’ (Domain 5) |
| 8: ALA143’ ARG134’ ASN137’ ASN144’ ASN152’ ASN161’ ASP143’ ASP155’ CYS148’ CYS153’ CYS157’ HIE150’ ILE158’ LEU139’ LEU140’ LEU147’ LYS136’ PHE146’ PRO138’ PRO159’ SER135’ PRO138’ PRO159’ SER135’ SER162’SER165’ SER166’ THR151’ THR164’ TRP149’ TYR141’ TYR145’ TYR154 TYR156’ TYR160’ VAL163’ (Domain 4) | 8: ALA143 ARG126 ARG134 ASN144 ASN152 ASN137 ASP142 ASP199 ASP210 ASP238 CYS148 CYS153 CYS157 CYS200 CYS130 CYS133 HIE150 HIE204 ILE158 ILE236 LEU139 LEU140 LEU147 LEU203 LEU127 LEU129 LYS198 LYS235 LYS132 LYS136 PHE146 PHE207 PRO138 SER205 SER209 SER135 THR151 THR208 TRP149 TRP128 TRP131 TYR141 TYR145 TYR154 TYR155 TYR156 TYR206 TYR211 | 8: ALA143 ARG122 ARG126 ARG134 ASN144 ASN161 ASN137 ASP142 ASP210 ASP238 CYS130 CYS133 HIE204 ILE236 ILE123 ILE124 LEU139 LEU140 LEU203 LEU127 LEU129 LYS235 LYS132 LYS136 MET125 PHE146 PHE207 PRO159 PRO138 SER162 SER205 SER209 SER135 THR208 TRP128 TRP131 TYR141 TYR145 TYR160 TYR206 TYR211 TYR212 VAL201 VAL202 VAL237 (Domains 4, 5) |

| Continuation of Table S4 | | |
|---|---|---|
| WT | D155Y | S171L |
| | TYR212 VAL201 VAL202 VAL237 (Domains 3, 4) | |
| 9: ARG134 ASN152 ASP155 ASP173 ASP183 CYS148 CYS153 CYS157 GLN185 GLU181 GLU191 GLU194 GLY172 GLY174 GLY187 GLY188 GLY196 HIE150 HIE182 ILE158 ILE179 ILE186 LEU147 LYS192 PRO178 SER177 SER180 SER195 SER135 THR151 THR175 THR176 THR190 TRP149 TRP193 TYR154 TYR156 TYR184 TYR189 VAL197 (Domain 4) | 9: ASP173 ASP183 GLN185 GLU181 GLU191 GLU194 GLY172 GLY174 GLY187 GLY188 GLY196 HIE182 ILE179 ILE186 LYS192 PRO178 SER177 SER180 SER195 THR175 THR176 THR190 TRP193 TYR184 TYR189 VAL197 (Domain 6) | 9: ASP173 ASP183 GLN185 GLU181 GLU191 GLY172 GLY174 GLY187 GLY188 HIE182 ILE179 ILE176 LEU171 PRO178 SER177 SER180 THR175 THR176 THR190 TYR184 TYR189 (Domain 6) |
| 10: ARG126 ASN137 ASP152 CYS130 CYS133 LEU139 LEU140 LEU127 LEU129 LYS132 LYS136 PRO138 TRP128 TRP131 TYR141 (Domain 4) | 10: ASN161 ASN234 ASP222 GLN213 GLN218 GLU226 GLY224 HIE227 ILE167 ILE169 ILE232 LEU214 LEU219 PHE230 PHE231 PRO159 SER162 SER165 SER166 SER171 SER216 SER220 THR164 THR170 THR217 THR221 THR223 THR229 TYR160 TYR215 TYR233 VAL163 VAL168 VAL225 VAL228 (Domains 5, 6) | 10: ASN152 ASP155 ASP199 CYS148 CYS153 CYS157 CYS200 GLU194 GLY196 HIE150 ILE158 LEU147 LYS192 LYS198 SER195 THR151 TRP149 TRP193 TYR154 TYR156 VAL197 |
| 11: ASN234 ASP222 GLN213 GLN218 GLU226 GLY224 HIE227 ILE167 ILE169 ILE232 LEU214 LEU219 PHE230 PHE231 SER171 SER216 SER220 THR170 THR217 THR221 THR223 THR229 TYR215 TYR233 VAL168 VAL225 VAL228 | 11: ASP173’ ASP183’ GLN185’ GLU181’ GLU191’ GLU194’ GLY172’ GLY174’ GLY187’ GLY188’ GLY196’ HIE182’ ILE179’ ILE186’ LYS192’ PRO178’ SER177’ SER180’ SER195’ THR175’ THR176’ THR190’ TRP193’ TYR184’ TYR189’ VAL197’ (Domain 6) | 11: ASN234 ASP222 GLN213 GLN218 GLU226 GLY224 HIE227 ILE167 ILE169 ILE232 LEU214 LEU219 PHE230 PHE231 SER165 SER166 SER216 SER220 THR164 THR170 THR217 THR221 THR223 THR229 TYR215 TYR233 VAL163 VAL168 VAL225 VAL228 |
| 12: ALA143 ASN144 ASN161 ASP199 ASP210 ASP238 CYS200 HIE204 ILE237 LEU203 LYS198 LYS235 PHE146 PHE207 PRO159 | 12: ASN234’ ASP222’GLN213’ GLN218’ GLU226’ GLY224’ HIE227’ ILE167’ ILE169’ ILE242’ LEU214’ LEU219 PHE230’ PHE231’ SER171’ | 12: ASP173’ ASP183’ GLN185’ GLU181’ GLU191’ GLU194’ GLY172’ GLY174’GLY187’ GLY188’ GLY196’ HIE182’ ILE179’ ILE186’ LEU171’ |

| Continuation of Table S4 | | |
|---|---|---|
| WT | D155Y | S171L |
| SER162 SER165 SER166 SER205 SER209 THR164 THR208 TYR145 TYR160 TYR206 TYR211 TYR212 VAL163 VAL201 VAL202 VAL237 (Domains 4, 5) | SER216’ SER220’ THR170’ THR217’ THR221’ THR223’ THR229’ TYR215’ TYR233’ VAL168’ VAL225’ VAL228’ | LYS192’ PRO178’ SER177’ SER180’ SER195’ THR175’ THR176’ THR190’ TRP193’ TYR184’ TYR189’ VAL197’ (Domain 6) |
| 13: ASP173’ ASP183’ GLN185’ GLU181’ GLU191’ GLU194’ GLY172’ GLY174’ GLY187’ GLY188’ GLY196’ HIE182’ ILE179’ ILE186’ LYS192’ PRO178’ SER177’ SER180’ SER195’ THR175’ THR176’ THR190’ TRP193’ TYR194’ TYR189’ VAL197’ (Domain 6) | | |
| 14: ASN234’ ASP222’ GLN213’ GLN218’ GLU226’ GLY224’ HIE227’ ILE167’ ILE169’ ILE242’ LEU214’ LEU219’ PHE230’ PHE231’ SER171’ SER216’ SER220’ THR170’ THR217’ THR221’ THR223’ THR229’ TYR215’ TYR233’ VAL168’ VAL225’ VAL228’ (Domain 6) | | |
| End of Table | | |

Table S5: **Rearrangement of constituent interactome clusters**. Every coloured box denotes a range of residues residing in the same cluster. Red bordered rectangles show how some of the residue sequences (coloured boxes) have merged in different combinations to form larger clusters due to increased strength of interactions between them.

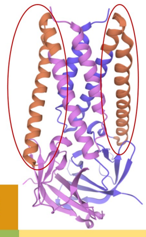

**Domain 3**

| WT | D155Y | S171L |
|---|---|---|
| 93-95 | 93-95 | 93-95 |
| 96-100 | 96-100 | 96-100 |
| 101-121 | 101-121 | 101-121 |
| 122-125 | 122-125 | 122-125 |
| 126-133 | 126-133 | 126-133 |
| 93’-95’ | 93’-95’ | 93’-95’ |
| 96’-117’ | 96’-117’ | 96’-117’ |
| 118’-125’ | 118’-125’ | 118’-125’ |
| 126’-133’ | 126’-133’ | 126’-133’ |

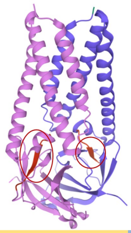

**Domain 4**

| WT | D155Y | S171L |
|---|---|---|
| 141’-149’ | 141’-149’ | 141’-149’ |
| 141-142 | 141-142 | 141-142 |
| 143-146 | 143-146 | 143-146 |
| 147-149 | 147-149 | 147-149 |

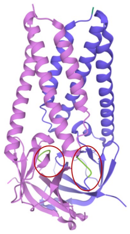

**Domain 5**

| WT | D155Y | S171L |
|---|---|---|
| 160’-163’ | 160’-163’ | 160’-163’ |
| 160-163 | 160-163 | 160-162 |
| | | 163 |

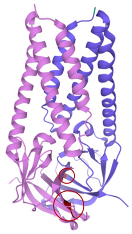

**Domain 6**

| WT | D155Y | S171L |
|---|---|---|
| 171’ | 171’ | 171’ |
| 172’-173’ | 172’-173’ | 172’-173’ |
| 171 | 171 | 171 |
| 172-173 | 172-173 | 172-173 |

Table S6: The list of PROVEAN scores of selected mutations of ORF3A.

| Amino acid Substitution in ORF3a | PROVEAN score* | Variation effect on protein |
|---|---|---|
| Q57H | -3.286 | Deleterious |
| W131C | -7.752 | Deleterious |
| W131R | -9.067 | Deleterious |
| D155Y | -6.829 | Deleterious |
| S171L | -2.238 | Neutral |
| G172C | -6.752 | Deleterious |
| G172V | -6.762 | Deleterious |

* Cutoff value = -2.5

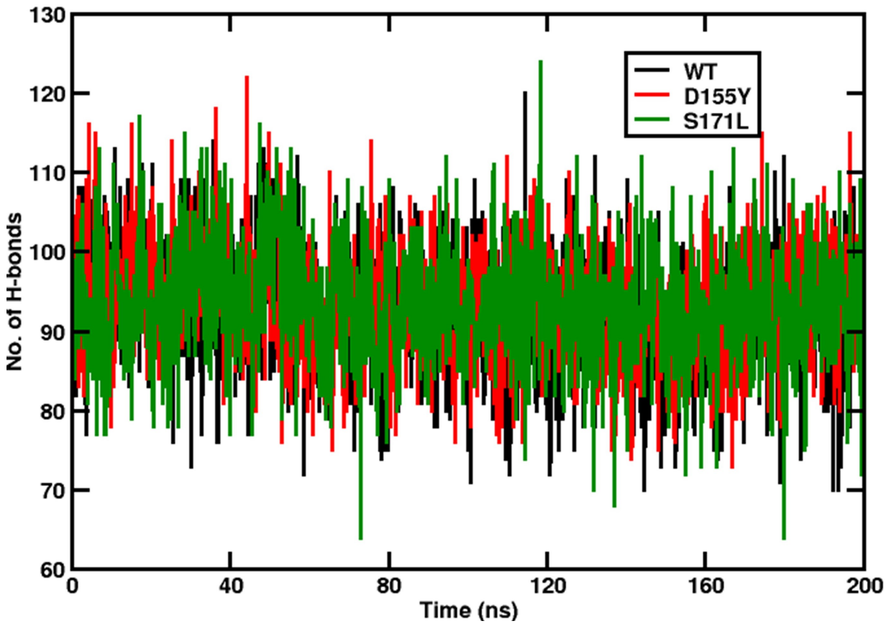


Fig.S1 The time evolution of hydrogen bonds for WT, D155Y and S171L are shown in black, red and green respectively

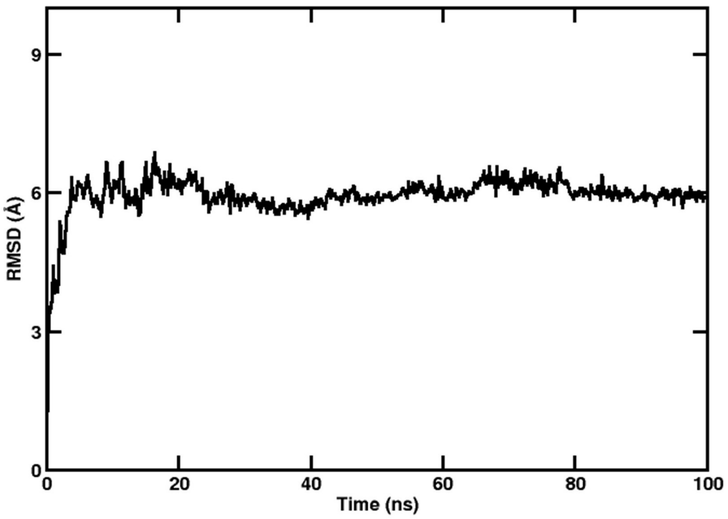


(a)

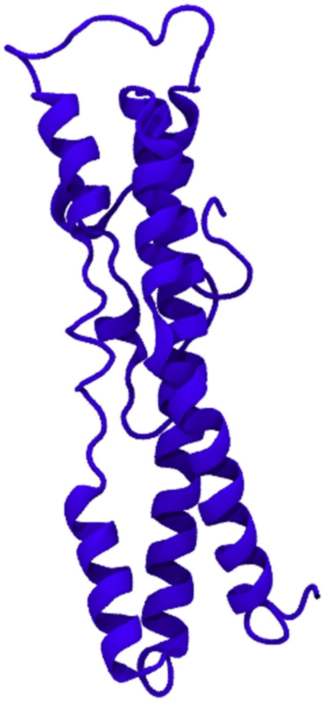

(b)

Fig. S2 (a) RMSD of the simulation of apo-caveolin-1 (b) Average structure of Caveolin-1 generated from simulation

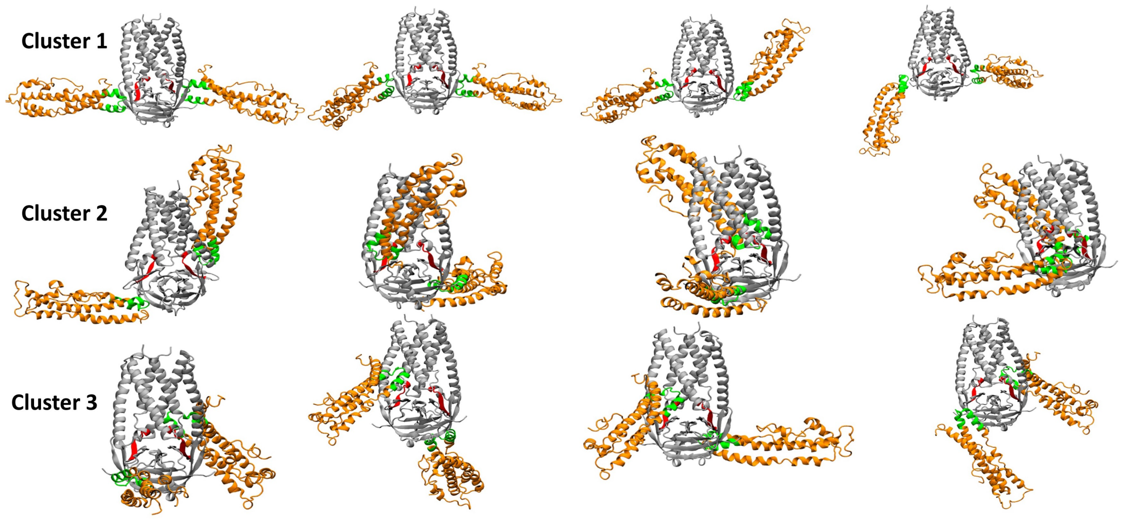


Fig. S3 The probable complexes generated from Haddock

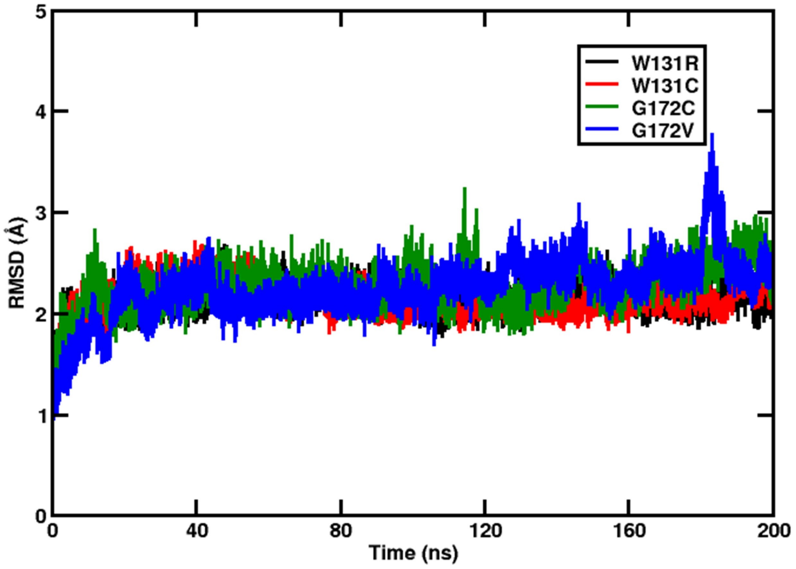


Fig. S4 The time evolution of RMSD of the four other mutants of ORF3a studied

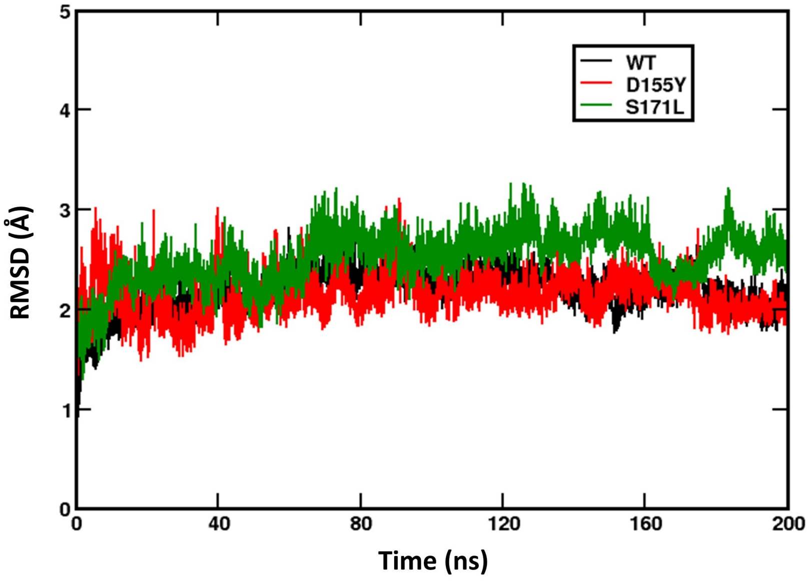


Fig. S5 The time evolution of the RMSD of the ORF3a protein with respect to the starting structure, obtained from the second production run

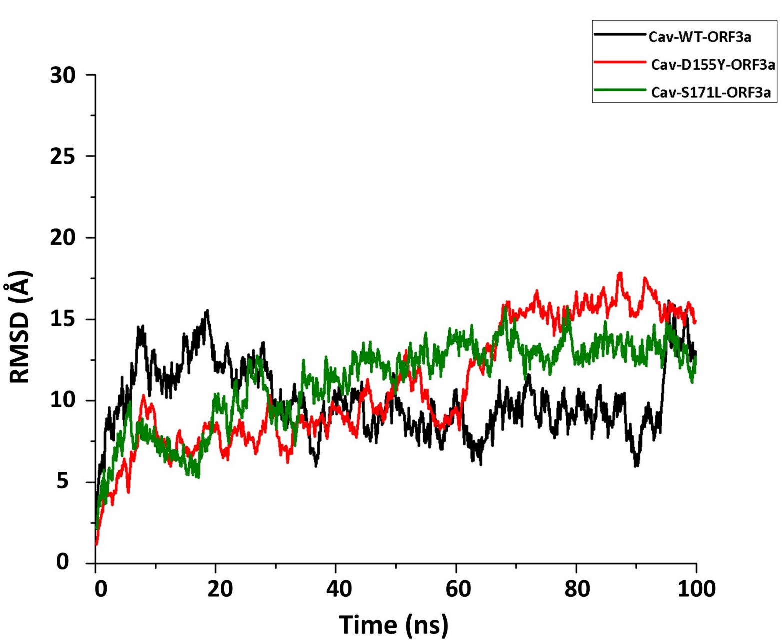


Fig. S6 Stability of the ORF3a-caveolin-1 complex, obtained from the second production run

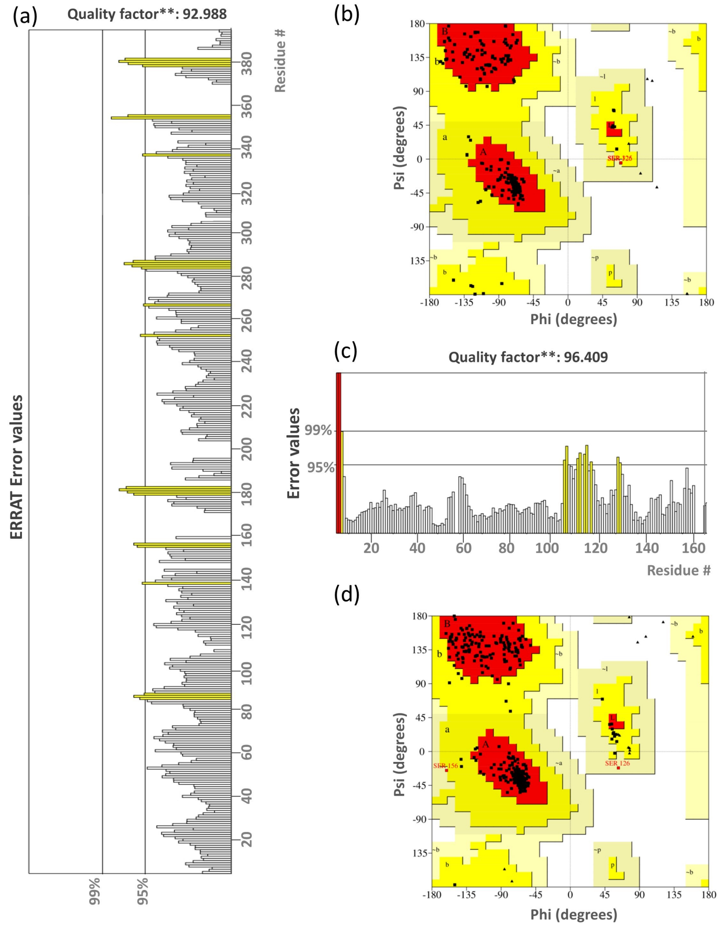


Fig. S7 The ERRAT analysis and Ramachandran Plot of ORF3a
